# Increased Relative Biological Effectiveness of Orthovoltage X-rays Compared to γ-rays in Preclinical Irradiation

**DOI:** 10.1101/2022.02.18.480594

**Authors:** Brett I. Bell, Justin Vercellino, N. Patrik Brodin, Christian Velten, Lalitha Sarad Yamini Nanduri, Kathryn E. Tanaka, Yanan Fang, Yanhua Wang, Rodney Macedo, Jeb English, Michelle M. Schumacher, Phaneendra K. Duddempudi, Patrik Asp, Wade Koba, Shahin Shajahan, Laibin Liu, Wolfgang Tomé, Weng-Lang Yang, Richard Kolesnick, Chandan Guha

**Affiliations:** Department of Radiation Oncology, Montefiore Medical Center, Bronx, NY, USA; Department of Pathology, Albert Einstein College of Medicine, Bronx, NY, USA; Department of Biochemistry, Albert Einstein College of Medicine, Bronx, NY, USA; Department of Radiology, Albert Einstein College of Medicine, Bronx, NY, USA; Laboratory of Signal Transduction, Memorial Sloan-Kettering Cancer Center, New York, NY 10065

**Keywords:** Radiobiology, Normal tissue response to radiation, Preclinical Models, Hematopoiesis, Immunology, Somatic Stem Cells

## Abstract

**Purpose:** Radionuclide irradiators (^137^Cs and ^60^Co) are commonly used in preclinical studies ranging from cancer therapy to stem cell biology. There are institutional initiatives to replace radionuclide sources with lower-energy X-ray sources amidst concerns of radiological terrorism. As researchers transition, there are questions whether the biological effects of γ-rays may be recapitulated with orthovoltage X-rays, since different energy may cause different biological effects. We, therefore, sought to compare the effects of orthovoltage X-rays and ^137^Cs γ-rays using mouse models of acute radiation syndrome.

**Experimental Design:** ^137^Cs γ-rays were compared with Orthovoltage X-rays, generated at 300 kVp, 10 mA with 1 mm Cu or Thoraeus filtration. We assessed 30-day overall survival following whole-body irradiation and calculated LD_50_ by logistic regression. Comparing equivalent doses delivered with different average energies (Ē), we assessed bone marrow, spleen, and intestinal histology and flow cytometry.

**Results:** The LD_50_ doses are 6.7 Gy, 7.4 Gy and 8.1 Gy with 1 mm Cu filtered (Ē=120 keV), and Thoraeus filtered X-rays (Ē=160 keV), and ^137^Cs (E=662 keV), respectively. At constant dose, hematopoietic injury was most severe with 1 mm Cu filtered X-rays with the greatest reduction in bone marrow cellularity, stem and progenitor populations, and intestinal crypts and OLFM4^+^ intestinal stem cells. Thoraeus filtered X-rays provoked an intermediate phenotype, with ^137^Cs showing the least damage.

**Conclusions:** Our study reveals a dichotomy between physical dose and biological effect relevant as researchers transition to orthovoltage X-rays. With decreasing energy, there is increasing hematopoietic and intestinal injury, necessitating dose-reduction to achieve comparable biological effects.

**Statement of Translational Relevance:** Radiation is used in translational studies in fields ranging from hematopoiesis and stem cell biology to cancer radiotherapy, with ^137^Cs and ^60^Co radionuclide sources serving as the most common irradiators. Due to the threat of radiological terrorism using stolen radionuclides, there are institutional initiatives to replace these sources with orthovoltage X-ray irradiators. Yet, as shown in this study, the biological effects of radiation are highly dependent on radiation energy. Lower energy orthovoltage X-rays are absorbed differently than higher energy radionuclide γ-rays, provoking more severe hematopoietic, immunologic, and gastrointestinal radiation injury. Thus, an identical physical dose delivered with beams of differing energy does not produce the same biologic effect. As researchers transition between these sources, it is critical that we appreciate that radiation doses are not interchangeable between them. Understanding the significance of physical dose delivered using different methods will allow us to contextualize past results with future studies.

## Introduction

Radiation is widely used as a tool for studies in cancer research, stem cell biology, and radiation countermeasures. Modern clinical irradiators consist predominantly of megavoltage (≥1 MV) linear accelerators or radionuclide γ-ray sources. High-energy radionuclide sources, such as ^137^Cs or ^60^Co, have been the predominant choice in preclinical studies, with only 30% of murine radiation studies reporting using lower energy orthovoltage (100-500 kVp) X-rays (1). This proportion, however, is rising steadily due to increased focus on the risk of theft and intentional dispersal of radionuclide sources. Therefore, The United States government has begun to encourage institutions to transition to X-ray sources to mitigate such risks (2). While replacing radionuclide sources with orthovoltage X-ray sources may ease security concerns, it represents a retreat to the past for radiobiology, the consequences of which must be carefully considered.

Orthovoltage X-ray therapy was once the dominant mode of treatment applied by early radiation oncologists. With the development of higher energy ^60^Co teletherapy units and megavoltage X-ray machines in the 1950s, the field shifted because such machines’ increased photon beam energies allowed for treatment of deeper tumors. This shift prompted studies that compared orthovoltage X-rays with the novel higher energy modalities and demonstrated significant differences in the biological response to radiation at the same physical dose depending on its energy (3). Extensive study of the relative biological effectiveness (RBE), a ratio of the doses of two types of radiation yielding the same effect on a biological system, subsequently quantified these differences. The effects of orthovoltage x-rays on cancer cells *in vitro* and murine tumors *in vivo* were remarkably more pronounced than radionuclide or megavoltage sources (4–6). The RBE between orthovoltage and megavoltage X-ray or radionuclide γ-ray sources was ultimately found to be clinically important enough to necessitate a 10% increase in dose to recapitulate the effects of orthovoltage cancer therapies with higher energy sources (7,8).

Early studies involving whole-body irradiation of mice found that comparing the radiation dose which causes 50% lethality from acute radiation syndrome (ARS) within 30-days (herein LD_50/30_ herein, referred to as LD_50_) was a useful measure to evaluate RBE (9–11). Although a few studies have suggested that orthovoltage sources produce comparable effects in certain preclinical applications (12,13), less attention has been paid recently to potential differences in biological effect, necessitating investigation with modern irradiators and techniques.

The different biologic effects seen with low and high energy ionizing radiation result from how radiation of different energies interacts with matter. At energies relevant to preclinical irradiation (∼100-1000 keV), two interactions of photons with matter dominate: photoelectric effect and Compton scattering. The probability of photoelectric absorption increases at low energies and with increasing atomic number of the interacting material, including bone. Compton scattering, in contrast, occurs more frequently with higher energy photons and is mostly independent of atomic number. Physical simulations and *in vitro* data have demonstrated that the mass dependence of photoelectric absorption increases the absorbed dose to bone marrow, especially with low energy photons (14).

Commercially available orthovoltage irradiators produce a spectrum of photon energies ranging from ∼100-350 peak kilovolts (kVp), with average energies approximately one-third of the peak, depending on the degree of filtration. Increased X-ray filtration may be used to reduce the fraction of low energy bremsstrahlung photons and characteristic X-rays, thereby increasing the average energy of the spectrum, potentially making orthovoltage sources behave more like higher energy radionuclide sources (1,14). Radionuclide γ-sources produce photons of discrete energies at 662 keV for ^137^Cs or 1173 keV and 1332 keV for ^60^Co. Given this range of energies, it is clear that preclinical irradiation walks the line between photoelectric and Compton absorption which may lead to diverse biological effects. We, therefore, aimed to assess for differential effects between orthovoltage X-rays, with varying degrees of filtration, and ^137^Cs γ-rays on survival, hematologic, immunologic, and gastrointestinal injury using models of acute radiation syndrome.

## Methods and Materials

### Animals

Eight to ten-week-old male C57BL/6J mice (IMSR Cat# JAX:000664, RRID: IMSR_JAX:000664) were randomized to groups on arrival and maintained in the animal facilities of the Albert Einstein College of Medicine. Mice were acclimated for one week prior to experiments and housed under specific pathogen free conditions with a 14:10 hour light:dark cycle (0600–2000), at 20-22°C and 30-70% humidity. Animals had *ad libitum* access to food (Lab Diet 5001) and water throughout the experiment. To limit pathogen transmission, water was acidified to a pH of 2.5-3.0 with HCl for survival studies. All experimental procedures were conducted in accordance with protocols approved by the Institutional Animal Care and Use Committee.

### Irradiation

For whole-body irradiation (WBI) using a Shepherd Mark I ^137^Cs irradiator, mice were anesthetized intraperitoneally with Ketamine/Xylazine and placed in a circular jig at the 100% isodose height on a turntable and irradiated according to manufacturer specifications at a dose rate ranging from 1.87-1.98 Gy/min due to radioactive decay of ^137^Cs over 2.5 years of study.

For X-ray WBI using a CIX-3 orthovoltage source (Xstrahl), un-anesthetized mice were placed into a plexiglass jig. The X-ray irradiator was operated at 300 kVp, 10 mA with either 1 mm Cu or Thoraeus (4 mm Cu Half-Value Layer (HVL)) filtration. For 1 mm Cu filtration, the dose rate was 1.89 Gy/min and for Thoraeus, 1.12 Gy/min, both at 40 cm source-surface distance. For partial-body irradiation (PBI) with X-rays using the above operating characteristics, unanesthetized mice were restrained in 50 mL conical tubes with the left lower limb exteriorized outside the tube to shield the tibia, fibula, ankle, and foot under lead. All irradiations were performed between 0900-1100h.

### Dosimetry

The manufacturer commissioned the CIX-3 orthovoltage irradiator using in-air measurements as previously described (15). Dose calculation protocols for small animal irradiation were established following acceptance testing of the irradiator and the dosimetric output was independently verified by the RNCP Irradiator Dosimetry Program. Thermoluminescent dosimeters (TLDs) calibrated for orthovoltage energy and traceable to NIST standards were exposed to a planned dose of 4.0 Gy. The average dose measured in two sets of TLDs by the independent laboratory were within 1.0% and 1.5% of the expected reading.

For quality assurance, experimentally delivered doses were verified using radiochromic film-based in-vivo dosimetry (16). A standard curve was generated by irradiating 2” x 2” pieces of GafChromic EBT-3 film, cut from the same lot and box, with 0 Gy to 15 Gy using a clinical linear accelerator’s 6 MV beam at the depth of maximum dose in solid water. Films were scanned in transmission mode using an EPSON Perfection V700. On the same day, film was irradiated using the CIX-3 or ^137^Cs. Doses were estimated from optical density using red and green channels in Matlab (MathWorks). All film was scanned in the same location using film holder. Due to the small film size and smaller regions of interest chosen for optical density measurement, no corrections for scanner position were performed. Film was irradiated using the CIX-3 and ^137^Cs to 5 Gy and 10 Gy doses in reference geometry to verify the consistency of the calibration curve for the CIX-3 under reference conditions. Film readout was always performed no earlier than 20 hours after irradiation and within 28 hours of irradiation. Continued development between 20 and 28 hours of irradiation was separately found to be negligible. Similarly, scanner heating was not found to impact film readings for few readings. Global changes in film optical density between different experiments were corrected via optical density shifts based on the reading from a piece of film irradiated under reference conditions on the CIX-3 immediately before or after the experiment.

### Health Status Monitoring and Survival

All animals were assessed daily, or twice daily during critical periods when death was expected, and weighed throughout the study. Published scoring criteria (MISS 3) were utilized to ensure consistent, humane euthanasia was completed when mice met pre-determined endpoints (17).

### LD_50_ Calculations

Dose-response models for 30-day mortality due to acute radiation syndrome were established based on the binary survival outcomes for animals irradiated using ^137^Cs, CIX-3 1mm Cu and CIX-3 Thoraeus, respectively. For each dose level the proportion of animals that died within 30 days was calculated along with binomial 95% confidence intervals. Probit dose-response models were fit to these data using the following formalism:

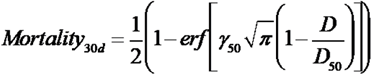

where *D_50_* represents the dose resulting in 50% mortality, *γ_50_* represents the normalized dose-response gradient and *erf* is the mathematical error function.

### Compton Scattering to Photoelectric Effect Ratio Calculation

Using NIST XCOM, we input values for the ICRU compact bone and ICRU four component soft tissue mixtures to determine the ratio of photoelectric effect and incoherent scattering in these materials at energies of 120 keV, 160 keV, and 662 keV.

### Complete Blood Count

Mice were euthanized by isoflurane overdose and blood was collected by cardiac puncture into K_2_EDTA coated tubes. Automated complete blood count with differential was performed using a Hemavet 950FS (Drew Scientific). Kwik-Diff-stained blood smears were evaluated for morphology.

### Histology

Formalin-fixed, paraffin-embedded sternum, thymus, spleen, stomach and intestine were sectioned at 5 µm and stained with Hematoxylin and Eosin (H&E) and Perl’s Prussian blue using standard protocols. Histology experiments were replicated at least twice using a minimum of 3 biological replicates per group. Coded samples were assessed by independent, board-certified pathologists blind to the code. Slides were imaged on the P250 Slide Scanner (3DHISTEC) using the 20x objective. Crypt microcolony assays were performed as previously described (18) and crypt depths were measured from the base of the crypt to the crypt-villus junction.

### Immunohistochemistry

Slides were re-hydrated through graded alcohols then 20 minutes of antigen retrieval was performed in a vegetable steamer. Slides were cooled to room temperature then washed with TBS containing 0.025% Triton X-100 (TBST), then peroxidase blocked with Bloxall for 10 minutes followed by 1 hour of blocking with 10% goat serum and avidin-biotin blocking. Sections were incubated with rabbit monoclonal antibodies against olfactomedin-4 [D6Y5A] (Cell Signaling Technology Cat# 39141, RRID:AB_2650511, 1:400), Transferrin Receptor [EPR20584] (Abcam Cat# ab214039, RRID:AB_2904534, 1:400), or Myeloperoxidase [EPR20257] (Abcam Cat# ab208670, RRID:AB_2864724, 1:1000) overnight at 4°C then rinsed with TBST prior to detection using an anti-rabbit avidin-biotin amplification kit (Vector Laboratories Cat# PK-6101, RRID:AB_2336820). Vector NovaRed, Vector SG, or DAB were used as chromogens and sections were counterstained with hematoxylin prior to dehydration and mounting.

### EdU Cell Proliferation Assay

20 mg/kg 5-ethynyl-2’-deoxyuridine (EdU; Sigma-Aldrich) in PBS was injected i.p. 2 hours prior to euthanasia. EdU was detected according to the manufacturer’s instructions (Thermo-Fisher). EdU^+^ cells/crypt were assessed by counting 50 intact crypts. Regenerating crypts were defined as crypts having ≥5 EdU^+^ cells. Fluorescent images were obtained on the P250 Slide Scanner (3DHISTEC) using the 20x objective with constant exposure time within each experiment.

### Flow Cytometry

Hematopoietic stem and progenitor cells (HSPCs) and splenic immune cells were stained with Zombie NIR fixable viability dye (Biolegend). Immune cells were Fc-blocked with rat anti-mouse CD16/32 [2.4G2] (Becton-Dickinson) for fifteen minutes. Extracellular staining was performed and acquired using an Aurora flow cytometer (Cytek Biosciences) and analyzed using FlowJo (FlowJo, RRID:SCR_008520) version #10.8.1 in conjunction with the uniform manifold approximation and projection plugin (UMAP RRID:SCR_018217) version #3.1) (19). For UMAP analysis, all samples were downsampled and concatenated to generate representative plots. Hematopoietic stem and progenitor cells were gated using a published strategy (20). Details are available in the supplemental methods.

### Bone Marrow Transplant

Bone marrow was prepared by flushing bones harvested from 2 six-week-old male CD45.1 mice (IMSR Cat# JAX:002014, RRID:IMSR_JAX:002014) and pooled. 10 million cells in PBS were injected i.v. via tail vein 24 hours post-irradiation, or PBS alone was injected.

### Statistical Analysis

Statistical analysis was performed using Prism (GraphPad Prism, RRID:SCR_002798) version #9.2.0. Survival was compared using Kaplan-Meier curves with log-rank. One-way analysis of variance (ANOVA) was utilized to compare three or more groups while two-way ANOVA was utilized for multiparametric flow cytometry, each using Tukey’s multiple comparisons test. Non-irradiated controls were excluded from statistics in HSPC flow cytometry and shown only as reference since the primary comparison was between irradiated groups. Power analysis was not performed. Results were considered statistically significant when p < 0.05 or the Bonferroni-corrected p-value. Data are presented as mean ± SEM.

### Data Availability

Data were generated by the authors and are available on request.

## Results

### Increased Relative Biological Effectiveness of Orthovoltage X-Rays for Whole-Body Irradiation

We sought to compare the effects of whole-body irradiation (WBI) using ^137^Cs γ-rays or orthovoltage X-rays on Hematopoietic-Acute Radiation Syndrome (H-ARS). We immediately noticed that there were discrepancies in the dose required to provoke lethality secondary to H-ARS between methods. This prompted a more detailed comparison of our machines and how their physical parameters could influence our biological systems. We also performed additional dosimetry validation to ensure that equivalent physical doses were delivered across methods.

The average energy (Ē) of our least filtered (1 mm Cu) X-ray spectrum is 120 keV. We can harden the beam to Ē=160 keV by attenuating low energy x-rays with a Thoraeus filter, an alloy equivalent to 4 mm Cu filtration (Communication with Xstrahl). ^137^Cs irradiation has the highest energy with a monoenergetic 662 keV beam. Radiation absorption varies with beam quality (approximately a function of Ē), particularly in the range of orthovoltage X-rays, with lower energy radiation increasing the proportion of photoelectric absorption relative to Compton absorption. Therefore, we calculated the ratio of Compton scattering to photoelectric absorption occurring in compact bone surrounding the bone marrow. The ratio of Photoelectric to Compton absorption is greatest with our lowest Ē beam, 1 mm Cu, and is reduced by hardening the beam with Thoraeus filtration while orthovoltage beams produce two orders of magnitude more photoelectric absorption than the ^137^Cs beam in both bone and soft tissue (Fig. 1A). Given these factors and simulation data suggesting increased absorbed dose to bone marrow due to photoelectric effect (14) we expected our lower energy orthovoltage beams to increase the severity of hematopoietic injury relative to ^137^Cs.

**Figure 1:**
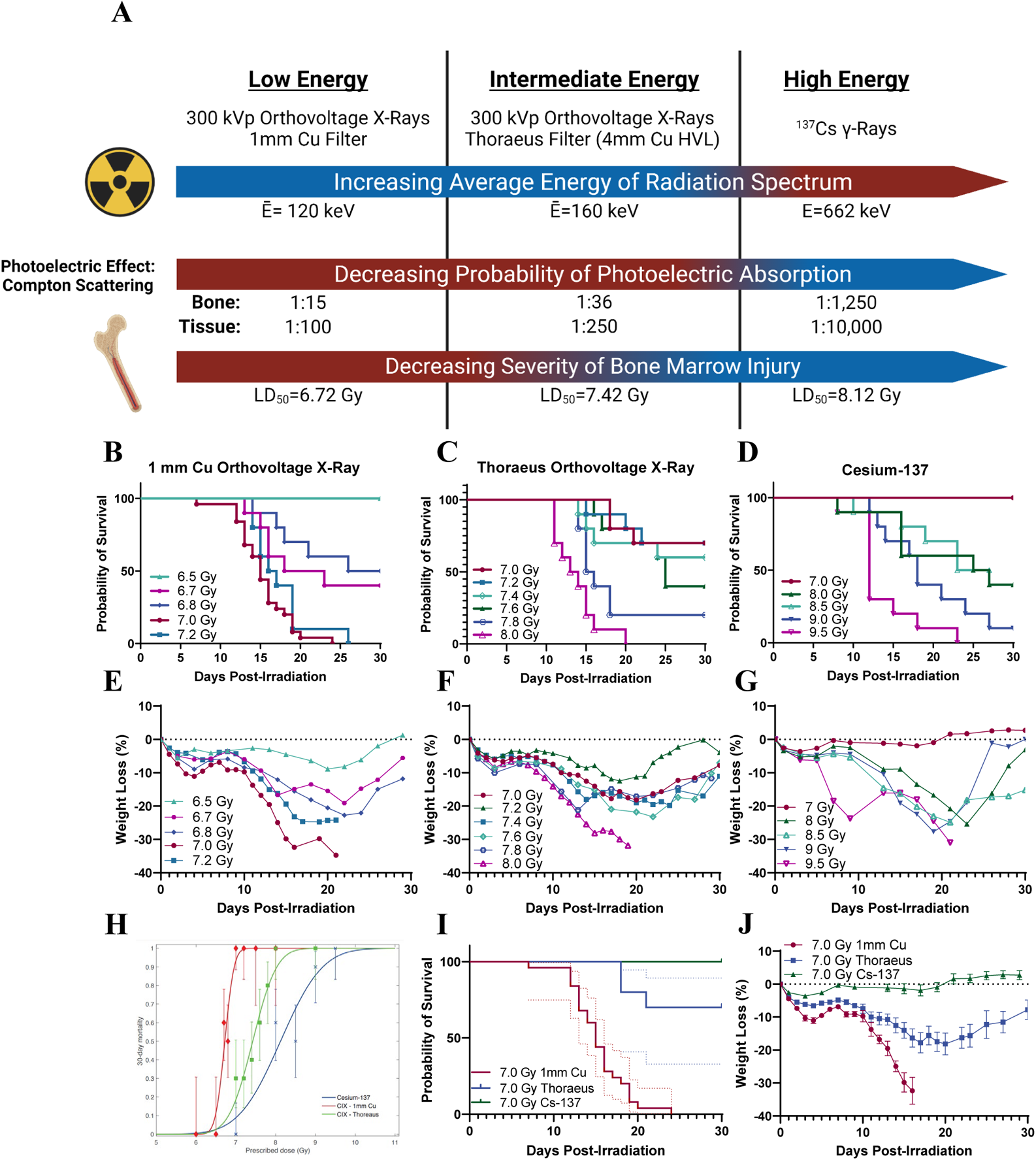
Radiation Beam Quality Alters 30-Day Survival in Mice Exposed to Whole-Body Irradiation. (A) Schematic depicting the effects of radiation energy on bone marrow injury with calculated average energies of the radiation spectra, ratios of photoelectric effect to Compton scattering in compact bone and soft tissue, and LD_50_ from each irradiation source. (B) Kaplan-Meier survival curves from experiments with whole-body irradiation of mice using orthovoltage X-rays with 1 mm Cu filtration, (C) orthovoltage X-rays with Thoraeus filtration, or (D) Cs-^137^ γ-rays. (n≥10/group; Results pooled from several experiments.) (E) Mean weight loss (%) following whole body irradiation with orthovoltage X-rays with 1 mm Cu filtration, (F) orthovoltage X-rays with Thoraeus filtration, or (G) Cs-^137^ γ-rays. (H) Logit analysis of 30-day survival using each radiation modality. (I) Kaplan-Meier curve comparing 7.0 Gy with each modality. 95% confidence intervals (dotted line). (J) Comparison of weight loss (%) following 7 Gy whole-body irradiation with each radiation modality. Data is represented as mean ± SEM. HVL: Half-Value Layer.

To test this hypothesis, we performed dose-response experiments to identify the lethal dose to provoke 50% mortality within 30-days (LD_50_) from H-ARS following whole-body irradiation (WBI; Fig. 1B-D). For 1 mm Cu filtered X-rays, the LD_50_ was 6.72 Gy (95% CI: 6.65, 6.80). Beam hardening with Thoraeus filtration increased the LD_50_ to 7.42 Gy (95% CI: 7.29, 7.55). ^137^Cs irradiation required the highest doses of radiation to induce death from H-ARS, with an LD_50_ of 8.12 Gy (95% CI: 7.44, 8.80). The RBEs for 1mm Cu and Thoraeus filtered X-rays relative to γ-rays were therefore 1.21 and 1.09, respectively. Similar trends were observed in weight loss, which increased with decreased radiation energy (Fig. 1E-G). Interestingly, 1 mm Cu demonstrated the steepest dose-response slope with ^137^Cs having the smallest slope, and Thoraeus intermediate (Fig 1H). Comparing 7.0 Gy irradiation highlights the differences in biological effect between groups with 100% lethality using 1 mm Cu filtered X-rays, reduced to 30% lethality with Thoraeus filtered X-rays (Thoraeus; *P* < 0.0001), and no lethality following ^137^Cs WBI (*P* < 0.0001; Fig. 1I). Weight loss was most severe post-IR with 1 mm Cu filtered X-rays and least severe following ^137^Cs irradiation, with Thoraeus filtered X-rays intermediate between the two (Fig. 1J). These experiments identified significant differences in survival depending on the quality of the radiation beam.

### Hematopoietic Injury Increases with Decreasing Radiation Energy Despite Delivering the Same Dose

Our previous experiments identified different doses necessary to induce the same effect on 50% lethality by 30-days. Here, we sought to assess the degree of hematopoietic injury at a constant dose of 7 Gy delivered by beams with differing radiation quality, given the distinct survival phenotypes observed. Preliminary data identified significant recovery from 7-14-days post-irradiation in ^137^Cs irradiated mice (Supplemental Fig. 1A). We therefore opted to perform more detailed assessment of the status of the hematopoietic system 10-days post-irradiation due to the high mortality with 1 mm Cu after this point. While there were not apparent differences between groups in peripheral blood due to a lag between bone marrow and peripheral recovery, pancytopenia in all groups confirms that irradiation induced H-ARS (Fig. 2A-C, Supplemental Fig. 1B).

**Figure 2:**
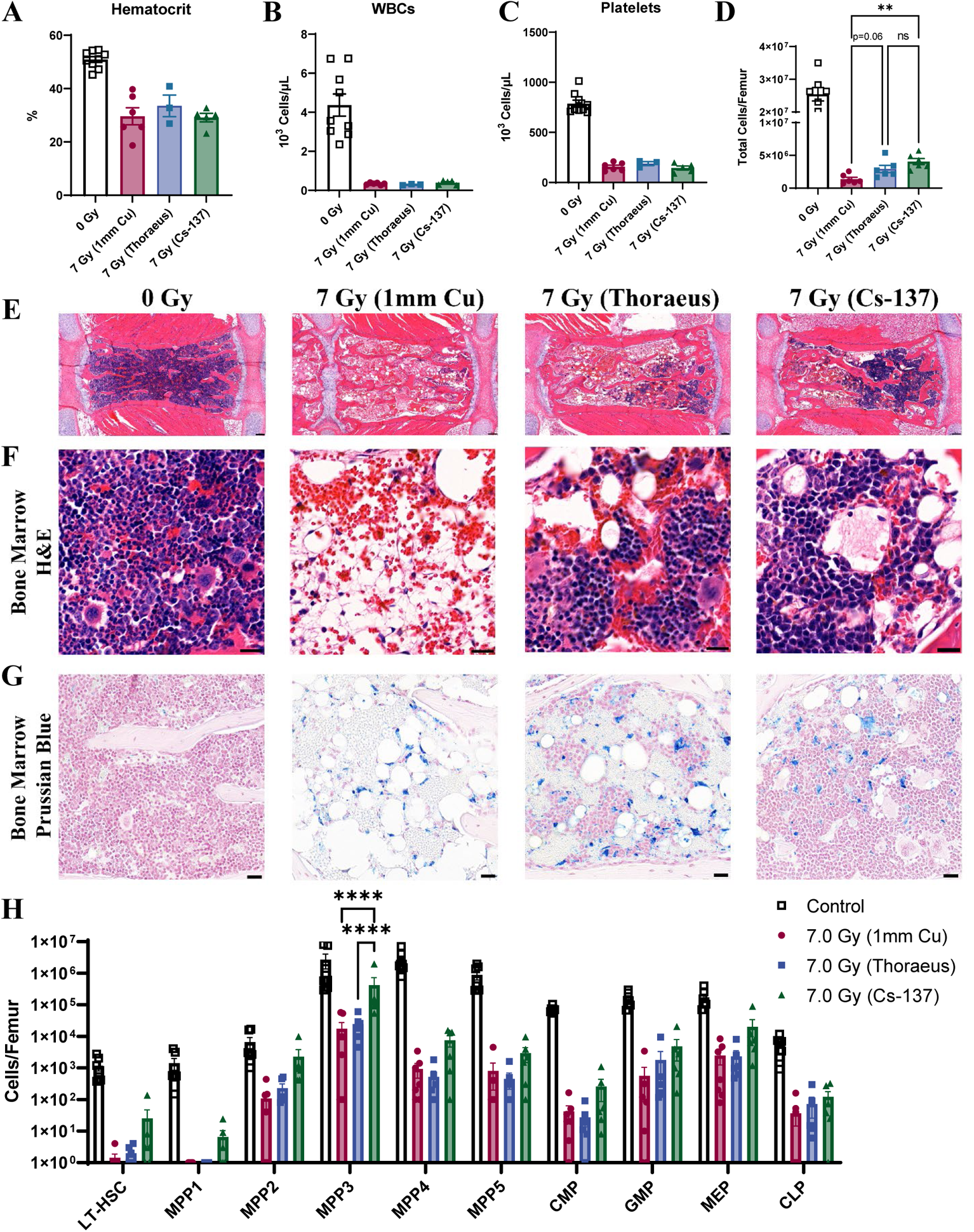
Hematopoietic Injury Decreases with Increasing Radiation Energy at a Constant Dose. Complete blood count analysis of peripheral blood for (A) hematocrit, (B) white blood cells, and (C) platelets 10-days after 7 Gy whole-body irradiation. Results combined from two independent experiments (n≥3/group). (D) Femoral cell counts were performed 10-days post-irradiation using an automated hemacytometer and combined from two independent experiments. (E) H&E-stained sternal bone marrow depicting sternebrae 10-days post-irradiation representative of three independent experiments. (Scale Bar 100 µm; n≥6/group). (F) H&E-stained sternal bone marrow 10-days post-irradiation representative of three independent experiments (Scale Bar 20 µm; n≥6/group). (G) Prussian blue staining with nuclear fast red counterstain to detect iron (blue) in sternal bone marrow 10-days post-irradiation. Representative of three independent experiments (Scale Bar 25 µm; n≥6/group). (H) Hematopoietic stem and progenitor cell flow cytometry was performed on bone marrow collected from mice 10-days post-irradiation with results combined from two independent experiments (n≥6/group). Populations are represented as the total number of cells per femur using logarithmic scale. Data is represented as mean ± SEM. LT-HSC: Long-Term Hematopoietic Stem Cell, MPP: Multipotential Progenitor, CMP: Common Myeloid Progenitor, GMP: Granulocyte-Macrophage Progenitor, MEP: Myeloid-Erythroid Progenitor, CLP: Common Lymphoid Progenitor.

10-days post-irradiation, all groups exhibited hypocellular marrow by femoral cell counts and sternal histology (Fig. 2D-E). ^137^Cs irradiated mice showed the most regeneration of the bone marrow with significantly increased cellularity (*P* = 0.0019) relative to 1 mm Cu treated mice, while Thoraeus filtration increased cellularity to a lesser degree (*P* = 0.0636). Both Thoraeus and ^137^Cs groups had focal regions of hematopoietic regeneration within sternal marrow. This contrasted with severely hypocellular marrow dominated by diffuse hemorrhage and increased adipose tissue in 1 mm Cu treated mice. Increased trilineage hematopoiesis was observed in Thoraeus and ^137^Cs irradiated mice, still with regions of hemorrhage and increased adiposity compared to control (Fig. 2E-F). Prussian blue staining identified increased intracellular and extracellular iron in irradiated groups over control bone marrow without apparent differences between groups (Fig. 2G; Supplemental Fig. 1C).

Flow cytometry for hematopoietic stem and progenitor cells (HSPCs) demonstrated a consistent trend where ^137^Cs irradiated mice had increased HSPCs of varying degrees of maturity compared to X-irradiated mice (Fig. 2H). Absolute HSPC counts from Thoraeus treated mice tended to lie either intermediate between those of 1 mm Cu and ^137^Cs treated mice or closer to those of 1 mm Cu treated mice. There were significant increases multipotent progenitor (MPP) 3 between ^137^Cs and 1 mm Cu or Thoraeus treated mice (*P* < 0.0001 each). These data demonstrate that the energy of irradiation alters the extent of myeloablation and hematopoietic reconstitution post-irradiation. Dose reduction is therefore necessary when transitioning between ^137^Cs and orthovoltage machines due to increased hematopoietic injury.

### Immunologic and Hematopoietic Recovery Varies with Radiation Energy Following Whole-Body Irradiation

We next aimed to assess the immunologic status of mice irradiated with beams of varying quality to determine whether this altered their immune recovery after administering the same physical dose of 7 Gy. Thymic atrophy with varying degrees of fibrosis was observed in the 1 mm Cu and Thoraeus treated groups 10-days post-irradiation compared to control. Meanwhile the ^137^Cs treated group exhibited reduced atrophy and fibrosis as well as more clearly delineated structure between the cortex and medulla (Fig. 3A). Thymic weights further suggested ^137^Cs treated mice recovered comparably to controls, with 1 mm Cu having the lowest weight and Thoraeus intermediate (Fig. 3B).

**Figure 3:**
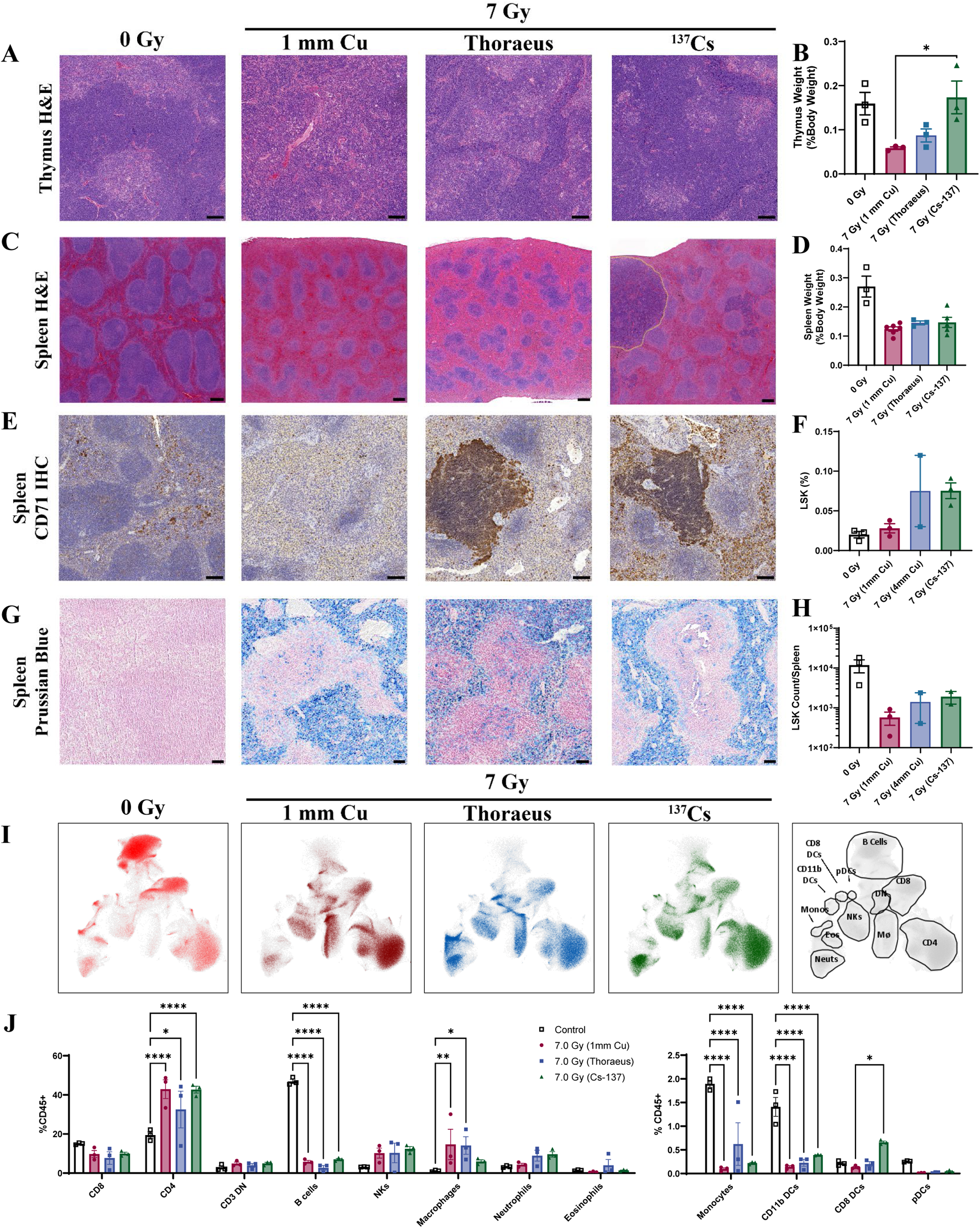
Splenic and Thymic Regeneration Following 7 Gy WBI Varies with Radiation Energy. (A) Representative images of H&E-stained thymus 10-days post-IR from two independent experiments (Scale Bar 100 µm; n≥3/group). (B) Thymus weights 10-days post-irradiation represented as a percent of body weight at the time of euthanasia. (C) H&E-stained spleens harvested from mice 10-days post-irradiation with regions of extramedullary hematopoiesis delineated by dotted yellow lines. Results are representative of two independent experiments. (Scale Bar 200 µm; n≥6/group). (D) Spleen weights 10-days post-IR represented as a percent of body weight at the time of euthanasia (n≥3/group). (E) CD71 IHC (brown) in spleens 10-days post-irradiation, representative of two independent experiments (Scale Bar 100 µm; n≥6/group). (F) Splenic flow cytometry for the percent of live Lin-, Sca-1+, c-Kit+ (LSK) cells 10-days post-irradiation (n≥2/group). (G) Spleens were stained with Prussian Blue to detect iron (blue) 10-days post-irradiation. Results are representative of three independent experiments. (H) Total number of LSK cells per spleen 10-days post-irradiation. (I) UMAP plots representing clusters of immune cell populations in the spleens 10-days post-irradiation. Gating strategy at right. (J) Leukocyte populations gated on UMAP clusters as a percent of total CD45^+^ cells. Data is represented as mean ± SEM. DN: Double Negative, NKs: Natural-Killer Cells, DCs: Dendritic Cells, pDCs: Plasmacytoid Dendritic Cells.

Splenic histopathology demonstrated atrophy of the white pulp with fibrosis in each irradiated group (Fig. 3C). The ^137^Cs irradiated group had increased white pulp with mildly expanded marginal zones compared to 1 mm Cu. This group additionally contained multifocal, often grossly visible, nodules of regenerating mononuclear cells in the red pulp most consistent with extramedullary hematopoiesis (EMH) 10-days post-irradiation, becoming more apparent by 14-days post-irradiation (Supplemental Fig. 2A). The Thoraeus group had fewer regenerative nodules while the 1 mm Cu treated group had none. There were no significant differences between groups concerning spleen weight (Fig. 3D).

To confirm that regenerative nodules represented EMH, we performed immunohistochemistry for transferrin receptor (CD71), a marker expressed on erythroid precursors, on spleens 10-days post-irradiation. These nodules were strongly CD71 positive in the ^137^Cs and Thoraeus treated groups, with the ^137^Cs group additionally showing increased CD71 positivity in the red pulp (Fig. 3E). Nodules were also myeloperoxidase positive, indicating myeloid differentiation (Supplemental Fig. 2B). HSPC flow cytometry confirmed increased percent and number of Lineage^-^, Sca-1^+^, c-Kit^+^ (LSK) cells in the Thoraeus and ^137^Cs groups, though individual comparisons did not rise to significance (Fig. 3F,H). This population contains all MPPs and is approximately 20% HSCs (21). These data suggest that EMH is increased in mice irradiated with higher energy sources.

Consistent with bone marrow H&E, we noted increased hemosiderin in irradiated spleens which led us to perform Prussian blue staining to assess iron status post-irradiation. Interestingly, there was severe iron overload in all irradiated spleens 10-days post-IR without apparent differences between irradiated groups (Fig. 3G) persisting until 14-days post-irradiation, while no hepatic increase was detected (Supplemental Fig. 2C-D).

After observing histologic differences in the spleen, we performed flow cytometry and visualized leukocyte populations via UMAP dimensional reduction (Fig 3I). We then validated our gating manually to identify clusters (Supplemental Fig. 3A). Irradiated groups generally experienced significant changes in immune populations relative to control. In terms of absolute number, radiosensitive lymphocyte populations decreased causing a relative increase in the percent of other populations post-irradiation (Supplemental Fig. 3B). This included increased percent CD4^+^ T-cells (1 mm Cu *P* < 0.0001, Thoraeus *P* = 0.0117, ^137^Cs *P* < 0.0001), and macrophages (1 mm Cu *P* = 0.0086, Thoraeus *P* = 0.0125, ^137^Cs *P* = 0.6367), as well as decreased percent B-cells (*P*<0.0001 each), monocytes (*P*<0.0001 each), and CD11b dendritic cells (DCs) (*P*<0.0001 each). While there were no significant differences comparing neutrophils between irradiated groups, there was a significant increase in percent CD8^+^ DCs in the ^137^Cs group relative to 1 mm Cu (*P* = 0.0304; Fig. 3J), revealing that the immunologic landscape differs depending on beam quality.

### Gastrointestinal Injury is Enhanced as Energy of Irradiation Decreases

Soft tissue injury as a function of radiation energy has not been well characterized. To address this, we assessed gastrointestinal injury, a significant component of ARS at higher doses. We evaluated the kinetics of gastrointestinal injury following 7 Gy WBI, finding only mild intestinal injury 4-days post-irradiation characterized by reactive crypt hyperplasia (Supplemental Fig. 4A). However, significant gastric hemorrhage was observed grossly 10-days post-IR and varied with radiation quality (Fig 4A-B*; P =* 0.0362). Gross samples were scored by a blinded observer who identified severe hemorrhage in the 1 mm Cu group and significant reductions in hemorrhage in the Thoraeus (*P* = 0.0288) and ^137^Cs groups (*P* = 0.0007). Histology confirmed hemorrhage largely within the body of the stomach in the 1 mm Cu group with prominent hemorrhage in the submucosa and in the lamina propria of the superficial mucosa (Fig. 4C).

**Figure 4:**
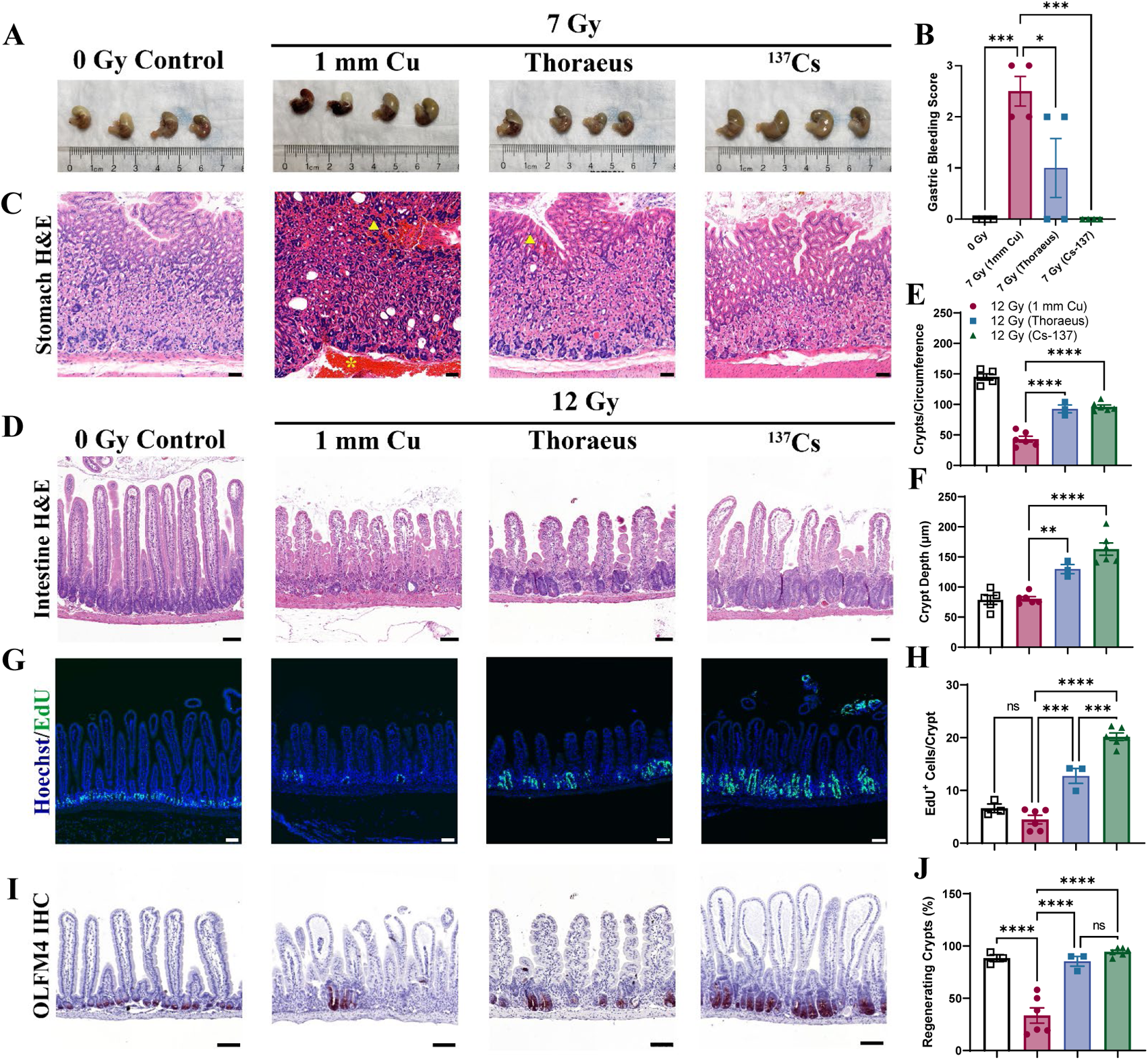
Gastrointestinal Injury is Dependent on Energy of Irradiation. (A) Gross pathology of stomachs 10-days after whole-body irradiation demonstrating gastric hemorrhage. (B) Gross gastric bleeding was scored semi-quantitatively by a blinded observer. (C) Representative H&E-stained sections of stomachs. Yellow arrowheads indicate mucosal hemorrhage. Yellow asterisk indicates submucosal hemorrhage. (Scale Bar 50 µm; n≥4/group). (D) H&E-stained intestinal sections from mice 4-days post-12 Gy WBI representative of two independent experiments. (Scale Bar 100 µm; n≥3/group.) (E) Total counts of crypts per circumference of intestinal cross-sections and (F) Crypt depth measurements 4-days post-irradiation. (G) EdU incorporation was fluorescently detected (green) to assess cell proliferation in intestinal crypts 4-days post-12 Gy WBI. Images representative from two independent experiments. (Scale Bar 100 µm; n≥3/group). (H) EdU^+^ cells per intestinal crypt were counted 4-days post-irradiation. (I) Olfactomedin-4 (OLFM4; red) was detected by IHC 4-days after 12 Gy of whole-body irradiation, representative of two independent experiments (Scale Bar 100 µm; n≥3/group). (J) The percent of regenerating crypts (≥5 EdU+ cells/crypt) was calculated 4-days post-irradiation. Data is represented as mean ± SEM.

Considering the relatively mild intestinal injury observed following 7 Gy WBI, we found it necessary to dose-escalate to elicit more substantial intestinal injury. We, therefore, irradiated mice with 12 Gy WBI, a dose often reported to deplete intestinal stem cells (ISCs) (22,23). 12 Gy WBI induced significant intestinal injury that varied with radiation quality 4-days post-irradiation in addition to provoking H-ARS with pancytopenia and hypocellular, hemorrhagic marrow (Supplemental Fig. 4B-C). Blinded histologic evaluation by an independent pathologist found that 1 mm Cu treated mice experienced the most severe intestinal injury by 4-days post-irradiation with significant recovery in both groups by day 7 (Fig. 4D, Supplemental Fig. 5A). There were many villi without crypts (crypt loss) or only vestigial crypts. The few crypts present showed minimal proliferative changes. Thoraeus treated mice had some crypt loss with most crypts exhibiting mild to moderate proliferative changes. ^137^Cs irradiated mice had some crypt loss with most crypts showing moderate proliferative changes.

Morphometric measurements confirmed these results, showing nearly a 60% reduction in crypts per cross-section in the 1 mm Cu group compared to control. Thoraeus and ^137^Cs caused significantly less crypt loss relative to 1 mm Cu (Fig. 4E; *P* < 0.0001 each). However, crypt depth did not increase in the 1 mm Cu group indicating a lack of proliferation in response to radiation. Crypt depths in the Thoraeus and ^137^Cs groups were significantly increased compared to the 1 mm Cu group (Fig. 4F; *P* = 0.0057 and *P* < 0.0001, respectively). ^137^Cs showed slightly increased crypt depth over Thoraeus (*P* = 0.0743). EdU staining detected minimal crypt epithelial cell proliferation in 1 mm Cu (Fig. 4G-H). ^137^Cs irradiated mice exhibited increased cell proliferation over 1 mm Cu (*P* < 0.0001) and Thoraeus (*P* = 0.0004), with Thoraeus also exhibiting increased proliferation above 1 mm Cu (*P* = 0.0001). 1 mm Cu however had a significant deficit of regenerating crypts (defined as ≥ 5 EDU^+^ cells/crypt; Fig. 4J; *P* < 0.0001) while Thoraeus and ^137^Cs increased to normal levels (*P* < 0.0001 for each).

Using immunohistochemistry to detect Olfactomedin-4 (OLFM4), a marker expressed on LGR5^+^ intestinal stem cells (ISCs) (24), we assessed ISC depletion post-irradiation. Following 7 Gy, there was no appreciable loss of ISC-containing crypts (Supplemental Fig. 5B). However, following 12 Gy 1 mm Cu, there was marked ISC loss relative to controls. Meanwhile, nearly all crypts in ^137^Cs treated animals contained OLFM4^+^ ISCs (Fig. 4I). These data demonstrate that intestinal injury and regeneration is dependent on the radiation source, with lower energy sources inducing more crypt and stem cell loss and slowing epithelial regeneration.

### Combined Hematopoietic and Gastrointestinal Injury Induced by Partial-Body Irradiation Depends on Radiation Quality

Partial-body irradiation (PBI) is used to study medical countermeasures for GI-ARS under the FDA animal rule (25–27). It intends to allow for dose-escalation to focus on GI-injury with the hematopoietic system partially intact. Using PBI with 2.5% bone marrow sparing by shielding the left-lower hindlimb, we assessed for differential survival after 1 mm Cu or Thoraeus treatment hoping to determine an RBE for intestinal injury. Here, we found that survival following 12 (*P* = 0.0050) or 13 Gy (*P* = 0.0021) was significantly improved, and weight loss was attenuated following PBI with Thoraeus versus 1 mm Cu, however there was limited dose-response which impaired RBE determination from survival data (Fig. 5A-D).

**Figure 5:**
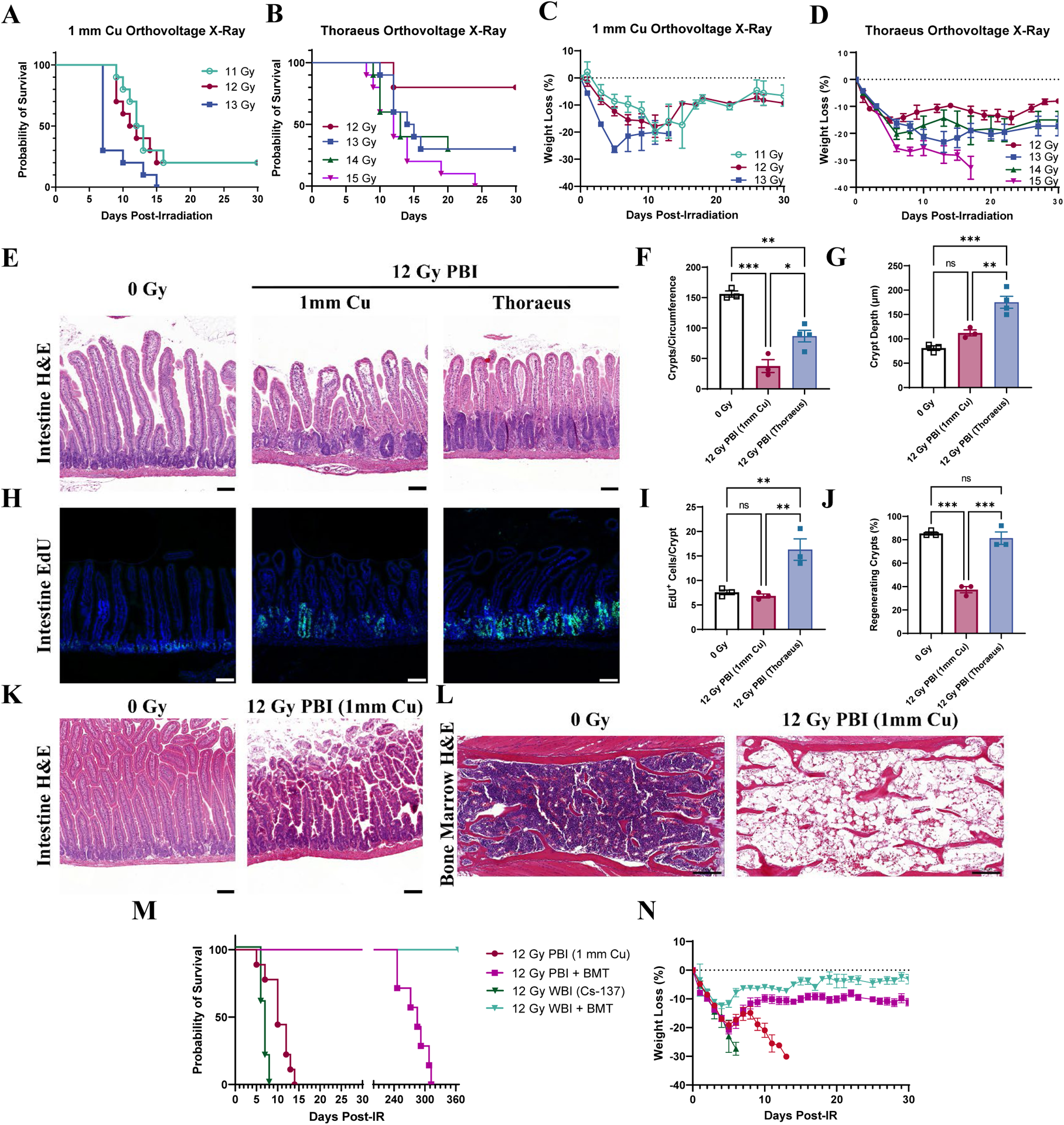
Response to Partial Body-Irradiation is Altered by Energy of Irradiation. (A) Kaplan-Meier survival curves for mice receiving partial-body irradiation with either 1 mm Cu or (B) Thoraeus filtered orthovoltage X-rays. (C) Comparison of weight loss (% of starting body weight) following partial-body irradiation with either (C) 1 mm Cu or (D) Thoraeus filtration. (E) Representative Hematoxylin & Eosin (H&E)-stained intestinal sections from mice 4-days post-12 Gy partial-body irradiation. (Scale Bar 100 µm; n≥3/group.) (F) Total counts of crypts per circumference of intestinal cross-sections and (G) Crypt depth measurements 4-days post-irradiation. (H) EdU incorporation was fluorescently detected (green) to assess cell proliferation in intestinal crypts 4-days post-12 Gy PBI (Scale Bar 100 µm; n≥3/group.) (I) EdU+ cells per intestinal crypt and (J) the percent of regenerating crypts (≥5 EdU+ cells/crypt) 4-days post-irradiation. (K) Representative H&E-stained intestinal sections from moribund mice following 12 Gy partial-body irradiation. (Scale Bar 100 µm; n=5). (L) Representative H&E-stained bone marrow sections showing one sternebrae in moribund mice (n=5). (M) Kaplan-Meier survival curves from mice irradiated either with 12 Gy PBI (1 mm Cu) or 12 Gy WBI (Cs- ^137^). Mice were either administered a PBS injection or bone marrow transplant 24 hours post-irradiation. (N) Weight loss as a percent of starting body weight for mice in the survival study. Data is represented as mean ± SEM.

We assessed small intestinal histopathology 4-days post 12 Gy PBI, a dose causing 80% lethality with 1 mm Cu but 20% lethality following Thoraeus filtered irradiation. Histopathologic differences in crypt depth, loss, and proliferation following 12 Gy PBI 1 mm Cu paralleled those seen after WBI (Fig. 5E). The degree of crypt loss was markedly less severe after 12 Gy PBI Thoraeus (*P =* 0.0141; Fig. 5F) with increased crypt depth 4-days post-irradiation over control (*P* = 0.0005) and 1 mm Cu irradiated animals (*P* = 0.0057; Fig 5G). After 12 Gy PBI, EdU^+^ cells increased in the Thoraeus group over both control and 1 mm Cu (*P* = 0.0078 and *P* = 0.0053; Fig. 5H-I). 1 mm Cu irradiation decreased the percent of regenerating crypts relative to control (*P* < 0.0002; Fig. 5J). Thoraeus treated mice also exhibited significant increases in regenerating crypts over 1 mm Cu treated animals (*P* = 0.0003; Fig. 5H-J) indicating a high degree of epithelial regeneration.

With 12 Gy being reported to ablate ISCs (23), we expected a significant contribution of GI-ARS to death at this dose. We carefully conducted necropsies on animals meeting euthanasia criteria post-12 Gy PBI 1 mm Cu to assess hematopoietic and gastrointestinal injury. Grossly, mice had hematomas and subcutaneous hemorrhages, pale livers and kidneys, as well as intestinal petechiae, all hallmarks of H-ARS. Intestinal histology at the time of death did not demonstrate damage consistent with death primarily from GI-ARS (Fig. 5K). Conversely, sternal marrow was aplastic with significant expansion of adipocyte populations and residual hemorrhage (Fig. 5L). The sum of these gross and histopathologic findings points towards H-ARS as the major contributor to death.

We then performed bone marrow transplants (BMT) to mitigate hematopoietic injury in an effort to understand gastrointestinal contributions to lethality post-irradiation. All mice receiving BMT survived, while those without died, suggesting that gastrointestinal injury alone was insufficient to cause death (Fig. 5M). The degree of weight loss in BMT treated mice was significantly attenuated with ^137^Cs compared to PBI 1 mm Cu, continuing to suggest an energy dependent biological effect (Fig. 5N).

We maintained the transplanted mice from this survival study for one-year post-irradiation to assess for delayed effects of acute radiation exposure. Mice treated with 12 Gy PBI 1 mm Cu + BMT began to die from delayed effects 8 months post-irradiation with 100% lethality by the 11^th^ month. Meanwhile, 100% of 12 Gy WBI ^137^Cs + BMT treated mice were alive one-year post-irradiation representing a significant difference (*P* = 0.0010) in delayed effects (Fig. 5M). These findings could be relevant while planning long-term hematopoietic reconstitution studies using radiation for myeloablation prior to BMT, with lower energy X-rays causing delayed death.

## Discussion

Researchers expect that the radiation dose administered will produce consistent biological effects regardless of radiation source. However, our work demonstrates that to produce the expected biological effect, care must be taken with respect to energy and filtration of the beam. While recent studies have demonstrated that orthovoltage X-rays and radionuclide γ-rays can achieve similar endpoints, including leukocyte depletion for blood transfusions clinically or myeloablation for BMT preclinically (12,13,28–30), orthovoltage sources are more biologically effective which can induce undesired toxicity when directly comparing doses (12).

30-day survival studies to determine LD_50_ were performed nearly 70 years ago to calculate the RBE of high energy irradiation versus orthovoltage X-rays, finding values ranging from 1.19-1.38 (Table 1). These RBEs were explained by increased photoelectric absorption in the bone producing secondary electrons, thus increasing absorbed dose in the bone marrow (31,32). Recent microdosimetry simulation and *in vitro* data demonstrates that increasing energy and filtration diminishes this effect (14). Our *in vivo* studies build on this data and are consistent with the notion that increasing X-ray filtration to increase the Ē of the radiation spectrum alters RBE. We observed remarkable concordance between our results and these prior studies, suggesting the effect is robust and reproducible (Table 1). The RBE of 1.21 using 1 mm Cu filtered X-rays indicates a ∼20% reduction in dose is necessary to produce the same biologic effect as ^137^Cs. Beam hardening with Thoraeus filtration shifts this by ∼10% to 1.09, requiring an only 10% reduction in dose with Thoraeus filtered X-rays. These values were within 4% and 1%, respectively, of calculated relative absorbed dose in bone marrow (14). This demonstrates that different quality beams have remarkably different effects on biologic systems and further proves that filtration may be used to narrow the gap in RBE between radionuclide and orthovoltage.

**Table 1:**
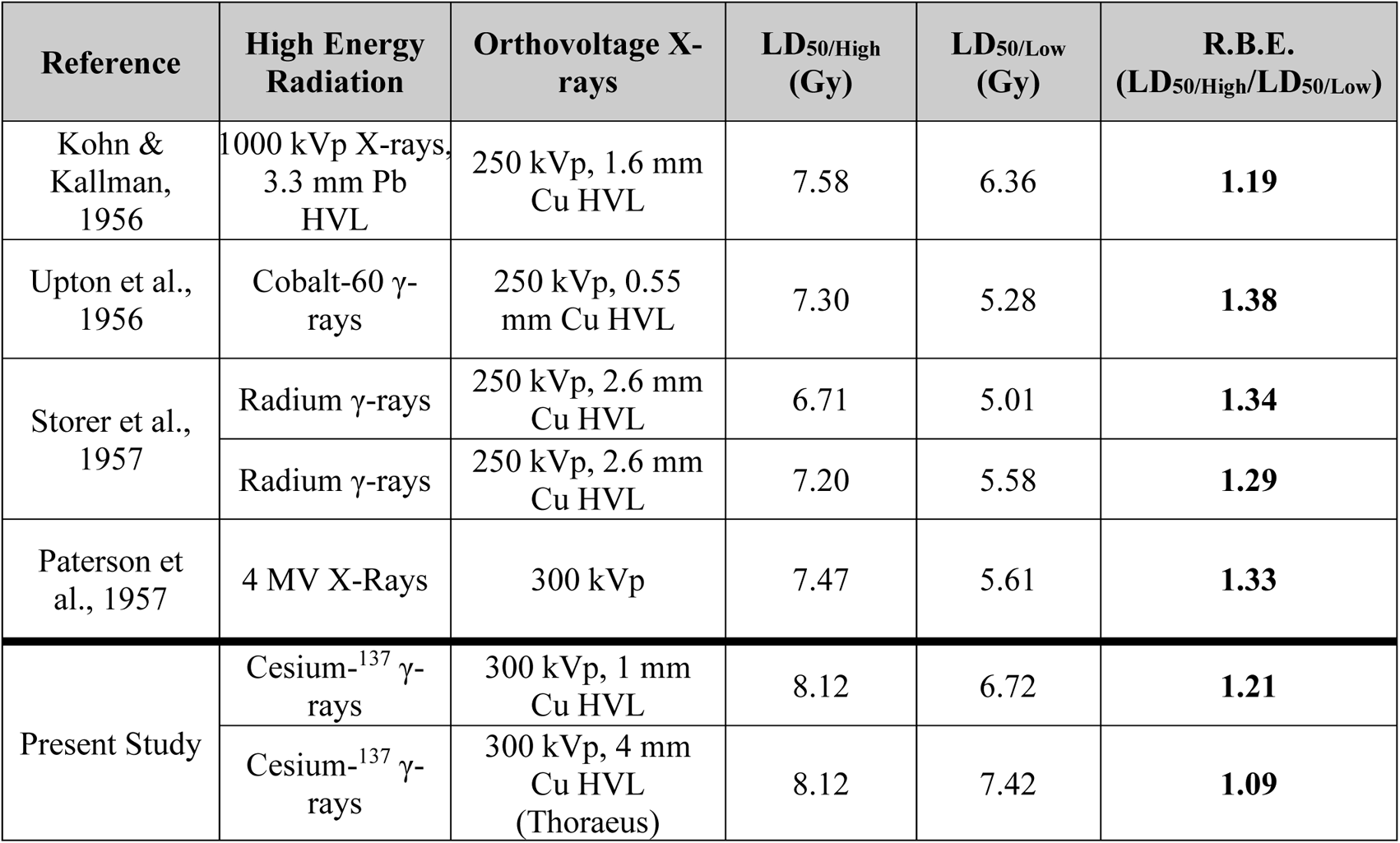
Comparison of Prior RBE Calculations with the Present Study. RBE was calculated using LD_50_ values from 30-day survival studies in mice after whole-body irradiation. Irradiation was performed with high energy radionuclide γ-ray sources or megavoltage X-ray sources and compared to orthovoltage X-rays. HVL: Half-Value Layer.

Multiple factors may influence RBE, however. Our orthovoltage WBI models utilized unanesthetized mice to better simulate radiation accidents or attacks, yet our longstanding ^137^Cs model utilized anesthetized mice. While the effects of anesthesia on WBI are debated (33–35), we believe that the difference in RBE between our un-anesthetized X-irradiated groups provides evidence that anesthesia effects were minor. Further, dose-rate may impact survival following WBI, although studies at lower dose-rates demonstrated that 30-day mortality from hematopoietic syndrome was only marginally affected by dose-rate (36). In our study, 1 mm Cu and ^137^Cs each exhibited dose-rates of ∼2 Gy/min allowing adequate comparison with constant dose-rate. Thoraeus filtered X-rays had a lower dose rate of 1.12 Gy/min, yet the expected intermediate RBE was identified, suggesting that dose-rate differences may have limited effect.

Bone marrow histology demonstrated that hematopoietic regeneration post-irradiation was dependent on radiation energy. Increased cellularity in Thoraeus and ^137^Cs irradiated bone marrow suggests a greater capacity for recovery, likely due to reduced marrow-ablation by these higher energy beams. This data correlated with flow cytometry demonstrating increases in HSPC populations, particularly in ^137^Cs irradiated mice, by 10-days post-irradiation. We anticipate that there would be greater separation between groups by 14-days since continued recovery increased marrow cellularity by this point, however ∼50% of 1 mm Cu irradiated mice are dead by then, presenting difficulties in studying these later timepoints. The differential effects seen in the marrow may not solely be due to HSPC loss but could also result from more severe stromal damage at lower energies which impairs hematopoietic regeneration and could also impact engraftment of donor HSPC after BMT (37). This is supported by modeling suggesting the increased absorbed dose in the marrow with lower energy X-rays is largely in the endosteal niche, which contains stromal populations responsible for supporting HSPCs (14,38).

In mice and humans, a significant proportion of hematopoiesis under stress conditions shifts to the spleen (39,40). We found that irradiation with 1 mm Cu did not induce splenic extramedullary hematopoiesis (EMH), while higher energies permitted development of EMH. Additionally, we evaluated primary and secondary lymphoid organs to understand immunologic recovery after irradiation with varying energies. Expansion of splenic white pulp and reduced thymic atrophy in the ^137^Cs group compared to 1 mm Cu speaks to a more rapid recovery after higher energy irradiation, with the Thoraeus group intermediate between the two.

Our observation that irradiation induced substantial iron accumulation within the bone marrow is consistent with prior work (41). Persistent iron overload within the marrow may generate reactive oxygen species by the Fenton reaction, causing a prolonged “second-hit” of oxidative stress, impairing hematopoietic recovery (42). Iron overload inhibits bone mineralization by osteoblasts and facilitates differentiation to osteoclasts, potentially contributing to bone deterioration post-irradiation (43,44). We observed severe splenic iron overload post-irradiation which could reflect splenic destruction of oxidized red blood cells to recycle iron. Radiation induces erythrocyte membrane alterations, including lipid peroxidation, that could prompt splenic recognition (45). This could partially account for the rapid onset of anemia (as early as 1-day post-irradiation) preceding gross signs of hemorrhage.

RBE is dependent on the biological system being studied and should be empirically determined for each system, therefore we attempted to calculate an intestinal RBE using survival data following PBI with 2.5% shielding of the marrow. However, we were unable to do so using our models as we have demonstrated doses sufficient to cause death from GI-ARS also induce concomitant H-ARS, because ARS is a multi-organ failure syndrome. Therefore, we must protect bone marrow to an extent such that intestinal injury is the primary contributor to death to determine an intestinal RBE using survival, such as, BMT after WBI or PBI.

Intestinal histologic recovery following 12 Gy PBI contrasted with aplastic bone marrow, and BMT studies demonstrated reconstitution of the marrow completely prevented death post-irradiation, suggesting H-ARS primarily contributed to death despite reports of LGR5^+^ ISC loss at this dose (22,23). This is consistent with prior work which demonstrates that 12 Gy WBI in C57BL/6J mice primarily leads to bone marrow failure, and dose-escalation is required to induce sufficient crypt loss to cause death primarily from intestinal failure (37,46–48). The influence of hematopoietic failure here despite the intent to achieve ISC depletion leads us to caution researchers against using physical dose as an absolute predictor of pathology. Instead, doses must be experimentally validated using individual irradiators to provoke the desired pathology. Furthermore, time-to-death should not be used as a sole indicator of the predominance of hematopoietic or gastrointestinal injury. While death in less than ten-to-fourteen days post-irradiation is frequently considered to be from GI-ARS, we observed this timing even at doses we are demonstrating to result from hematopoietic failure, particularly at lower energy, consistent with prior literature (37).

While we were unable to calculate an intestinal RBE based on overall survival in this study, one study calculated an RBE of 1.16 using survival of clonogenic crypt cells per circumference between 300 kVp X-rays (unknown filtration) and ^60^Co γ-rays (49). This indicates that histopathology may be useful in determining RBEs for intestinal radiation studies. Although additional study is needed, for which we suggest utilizing intestinal histopathology and functional endpoints in addition to survival. Comparing intestinal injury histologically at identical doses, while varying beam quality, revealed significant differences in crypt loss, stem cell ablation and epithelial proliferation between irradiated groups after both WBI and PBI. While there is good evidence that photoelectric absorption is increased in bone with orthovoltage X-rays due to the energy and atomic number dependence of the photoelectric effect (1,14), it is unclear if there is a similar physical influence which could increase absorbed dose in the intestine. The more severe intestinal injury we identified was accordingly unexpected, and further study is required to explain this phenomenon. It is possible, although unlikely, that the increased probability of photoelectric absorption could increase absorbed dose, as in bone. Increased density of ionization with lower energy X-rays could also contribute (50). In addition to these direct effects on soft tissue, the more severe hematopoietic injury following orthovoltage irradiation could hamper intestinal regeneration due to a role for bone marrow-derived cells in GI-ARS (51). Comparing between WBI and PBI with orthovoltage X-rays demonstrates improved histologic structure and morphometry by 4-days following PBI, supporting this notion of hematopoietic contributions to gastrointestinal recovery.

Consequently, we recommend against utilizing the RBE values calculated herein for dose-adjustment in the intestinal compartment because the measurement of RBE is inherently influenced by the specific biologic and physical parameters of the system.

Overall, we have demonstrated that within the range of energies used in preclinical irradiation, there is remarkable variation in biological effectiveness. Orthovoltage X-rays produced more severe hematopoietic, immunologic, and gastrointestinal injury compared to ^137^Cs γ-rays at constant dose. While our results are specific to these systems, the overarching concept transcends systems (3–11,49,52), and is therefore relevant to all researchers using radiation in their studies. Reporting a dose is insufficient without detailed reporting of source, energy, and filtration, preferably with characterization of biological effect. Knowledge of this phenomenon will aid in the interpretation and reproducibility of studies as researchers transition from radionuclide to orthovoltage sources.

As radiation countermeasures researchers, we cannot abide the continued use of radionuclide sources outside of specialized laboratories. While we have drawn attention to the difference in RBE between these sources, orthovoltage can be utilized in many, if not most, cases. This transition, however, necessitates physical dose-reduction and optimization.

## Author Contributions

Conceptualization, B.I.B., R.K., and C.G.; Methodology, B.I.B, J.V., N.P.B., C.V., and L.S.Y.N.; Investigation, B.I.B., J.V., L.S.Y.N., J.E., P.K.D., P.A., W.K., S.S., L.L.; Formal Analysis, B.I.B., J.V., N.P.B., C.V., K.E.T., Y.F., Y.W., R.M.; Writing – Original Draft, B.I.B.; Writing – Review & Editing; B.I.B., J.V., N.P.B., C.V., L.S.Y.N., K.E.T., Y.F., Y.W., R.M.G., J.E., M.M.S., P.K.D., P.A., W.T, W.Y., R.K., and C.G.; Visualization, B.I.B, J.V., N.P.B., R.G., and M.M.S.; Resources, C.G.; Funding Acquisition, C.G.; Supervision, W.T., W.Y., R.K., and C.G.

## References

1. Poirier Y, Belley MD, Dewhirst MW, Yoshizumi TT, Down JD, Poirier Y, et al. Transitioning from Gamma Rays to X Rays for Comparable Biomedical Research Irradiations: Energy Matters. RADIATION RESEARCH [Internet]. 2020 [cited 2022 Jan 5];193:506–11. Available from: http://meridian.allenpress.com/radiation-research/article-pdf/193/6/506/2512429/i0033-7587-193-6-506.pdf

2. Nuclear Regulatory Commission. The 2018 Radiation Source Protection and Security Task Force Report. Washington, D.C.; 2018.

3. Hall EricJ. The Relative Biological Efficiency of X Rays Generated at 220 kVp and Gamma Radiation from a Cobalt 60 Therapy Unit. http://dx.doi.org/101259/0007-1285-34-401-313 [Internet]. The British Institute of Radiology ; 2014 [cited 2022 Jan 5];34:313–7. Available from: https://www.birpublications.org/doi/abs/10.1259/0007-1285-34-401-313

4. Ting TP, Johns HE, Jaques LB. Relative Biological Effectiveness of Betatron and Conventional X-Radiation on the Regression of Mouse Tumours. Nature 1952 170:4331 [Internet]. Nature Publishing Group; 1952 [cited 2022 Jan 30];170:752–3. Available from: https://www.nature.com/articles/170752a0

5. Sugiura K. The Biological Measurement of Gamma Rays in “Equivalent Roentgens” with Mouse Sarcoma 180 as the Test Object. The American Journal of Cancer [Internet]. American Association for Cancer Research Journals; 1939 [cited 2022 Jan 30];37:445–52. Available from: https://cancerres.aacrjournals.org/content/37/3/445

6. Lasnitzki I, Lea DE. The Variation with Wavelength of the Biological Effect of Radiation (Measured by the Inhibition of Division in Tissue Cultures). http://dx.doi.org/101259/0007-1285-13-149-149 [Internet]. The British Institute of Radiology ; 2014 [cited 2022 Jan 30];13:149–62. Available from: https://www.birpublications.org/doi/abs/10.1259/0007-1285-13-149-149

7. Borek Carmia, Hall EricJ, Zaider Marco. X rays may be twice as potent as gamma rays for malignant transformation at low doses. Nature [Internet]. 1983 [cited 2022 Jan 5];301:156–8. Available from: 10.1038/301156a0

8. Amols HI, Lagueux B, Cagna D, Radiobiological D. Radiobiological Effectiveness (RBE) of Megavoltage X-Ray and Electron Beams in Radiotherapy. RADIATION RESEARCH [Internet]. 1986 [cited 2022 Jan 5];105:58–67. Available from: http://meridian.allenpress.com/radiation-research/article-pdf/105/1/58/2119411/3576725.pdf

9. Kohn HI, Kallman RF. Relative Biological Efficiency of 1000-Kvp and 250-Kvcp X-Rays: III. Determinations Based on the LD50/28 and the Killing Time of the Mouse. Source: Radiation Research. 1956;5:693–9.

10. Upton AC, Conte FP, Hurst GS, Mills WA. The relative biological effectiveness of fast neutrons, x-rays, and gamma-rays for acute lethality in mice. Radiation research. 1956;4:117–31.

11. Paterson E, Ashton M, Shaw J, Mayo J. The Relative Biological Efficiency of 4 MeV and 300 kV Radiations: A Symposium: IV. Experiments on Organ Weight Loss and 50 Per Cent Mortality in Mice. http://dx.doi.org/101259/0007-1285-30-355-343 [Internet]. The British Institute of Radiology ; 1957 [cited 2022 Jan 6];30:343–7. Available from: https://www.birpublications.org/doi/abs/10.1259/0007-1285-30-355-343

12. Gibson BW, Boles NC, Souroullas GP, Herron AJ, Fraley JK, Schwiebert RS, et al. Comparison of Cesium-137 and X-ray Irradiators by Using Bone Marrow Transplant Reconstitution in C57BL/6J Mice. American Association for Laboratory Animal Science;

13. Eng J, Orf J, Perez K, Sawant D, DeVoss J. Generation of bone marrow chimeras using X-ray irradiation: comparison to cesium irradiation and use in immunotherapy. Journal of Biological Methods [Internet]. Journal of Biological Methods; 2020 [cited 2022 Jan 17];7:e125. Available from: https://jbmethods.org/jbm/article/view/314

14. Belley MD, Ashcraft KA, Lee CT, Cornwall-Brady MR, Chen JJ, Gunasingha R, et al. Microdosimetric and Biological Effects of Photon Irradiation at Different Energies in Bone Marrow. https://doi.org/101667/RR140951 [Internet]. Radiation Research Society; 2015 [cited 2022 Jan 7];184:378–91. Available from: https://bioone.org/journals/radiation-research/volume-184/issue-4/RR14095.1/Microdosimetric-and-Biological-Effects-of-Photon-Irradiation-at-Different-Energies/10.1667/RR14095.1.full

15. Ma CM, Coffey CW, DeWerd LA, Liu C, Nath R, Seltzer SM, et al. AAPM protocol for 40-300 kV x-ray beam dosimetry in radiotherapy and radiobiology. Medical physics [Internet]. Med Phys; 2001 [cited 2022 Jan 30];28:868–93. Available from: https://pubmed.ncbi.nlm.nih.gov/11439485/

16. Niroomand-Rad A, Chiu-Tsao ST, Grams MP, Lewis DF, Soares CG, van Battum LJ, et al. Report of AAPM Task Group 235 Radiochromic Film Dosimetry: An Update to TG-55. Medical Physics [Internet]. John Wiley & Sons, Ltd; 2020 [cited 2022 Jan 30];47:5986–6025. Available from: https://onlinelibrary.wiley.com/doi/full/10.1002/mp.14497

17. Koch A, Gulani J, King G, Hieber K, Chappel M, Ossetrova N. Establishment of Early Endpoints in Mouse Total-Body Irradiation Model. PLOS ONE [Internet]. Public Library of Science; 2016 [cited 2022 Jan 17];11:e0161079. Available from: https://journals.plos.org/plosone/article?id=10.1371/journal.pone.0161079

18. Withers HR, Elkind MM. Microcolony Survival Assay for Cells of Mouse Intestinal Mucosa Exposed to Radiation. http://dx.doi.org/101080/09553007014550291 [Internet]. Taylor & Francis; 2009 [cited 2022 Feb 17];17:261–7. Available from: https://www.tandfonline.com/doi/abs/10.1080/09553007014550291

19. McInnes L, Healy J, Melville J. UMAP: Uniform Manifold Approximation and Projection for Dimension Reduction. 2018 [cited 2022 Jan 17]; Available from: https://arxiv.org/abs/1802.03426v3

20. Eich M, Trumpp A, Schmitt S. OMIP-059: Identification of Mouse Hematopoietic Stem and Progenitor Cells with Simultaneous Detection of CD45.1/2 and Controllable Green Fluorescent Protein Expression by a Single Staining Panel. Cytometry Part A [Internet]. John Wiley & Sons, Ltd; 2019 [cited 2022 Jan 19];95:1049–52. Available from: https://onlinelibrary.wiley.com/doi/full/10.1002/cyto.a.23845

21. Kiel MJ, Yilmaz ÖH, Iwashita T, Yilmaz OH, Terhorst C, Morrison SJ. SLAM Family Receptors Distinguish Hematopoietic Stem and Progenitor Cells and Reveal Endothelial Niches for Stem Cells. Cell. Cell Press; 2005;121:1109–21.

22. Asfaha S, Hayakawa Y, Muley A, Stokes S, Graham TA, Ericksen RE, et al. Krt19(+)/Lgr5(−) cells are radioresistant cancer initiating stem cells in the colon and intestine. Cell stem cell [Internet]. NIH Public Access; 2015 [cited 2022 Feb 3];16:627. Available from: /pmc/articles/PMC4457942/

23. Yan KS, Chia LA, Li X, Ootani A, Su J, Lee JY, et al. The intestinal stem cell markers Bmi1 and Lgr5 identify two functionally distinct populations. Proceedings of the National Academy of Sciences of the United States of America [Internet]. National Academy of Sciences; 2012 [cited 2022 Feb 12];109:466–71. Available from: https://www.pnas.org/content/109/2/466

24. van der Flier LG, van Gijn ME, Hatzis P, Kujala P, Haegebarth A, Stange DE, et al. Transcription Factor Achaete Scute-Like 2 Controls Intestinal Stem Cell Fate. Cell. Cell Press; 2009;136:903–12.

25. Fish BL, MacVittie TJ, Gao F, Narayanan J, Gasperetti T, Scholler D, et al. Rat Models of Partial-body Irradiation with Bone Marrow-sparing (Leg-out PBI) Designed for FDA Approval of Countermeasures for Mitigation of Acute and Delayed Injuries by Radiation. Health physics [Internet]. NLM (Medline); 2021 [cited 2022 Feb 12];121:419–33. Available from: https://journals.lww.com/health-physics/Fulltext/2021/10000/Rat_Models_of_Partial_body_Irradiation_with_Bone.11.aspx

26. Booth C, Tudor G, Tudor J, Katz BP, MacVittie TJ. Acute gastrointestinal syndrome in high-dose irradiated mice. Health physics [Internet]. Health Phys; 2012 [cited 2022 Feb 13];103:383–99. Available from: https://pubmed.ncbi.nlm.nih.gov/23091876/

27. MacVittie TJ, Farese AM, Parker GA, Jackson W, Booth C, Tudor GL, et al. The Gastrointestinal Subsyndrome of the Acute Radiation Syndrome in Rhesus Macaques: A Systematic Review of the Lethal Dose-response Relationship With and Without Medical Management. Health physics [Internet]. Health Phys; 2019 [cited 2022 Feb 13];116:305–38. Available from: https://pubmed.ncbi.nlm.nih.gov/30624353/

28. Andersen AHF, Nielsen SSF, Olesen R, Harslund JLF, Søgaard OS, Østergaard L, et al. Comparable human reconstitution following Cesium-137 versus X-ray irradiation preconditioning in immunodeficient NOG mice. PLOS ONE [Internet]. Public Library of Science; 2020 [cited 2022 Jan 30];15:e0241375. Available from: https://journals.plos.org/plosone/article?id=10.1371/journal.pone.0241375

29. Mackenzie C, Iwamoto KS, Smith K. University of California Replacement of Cesium Irradiators with Alternative Technologies. Health Physics [Internet]. Lippincott Williams and Wilkins; 2020 [cited 2022 Jan 30];118:209–14. Available from: https://journals.lww.com/health-physics/Fulltext/2020/02000/University_of_California_Replacement_of_Cesium.9.aspx

30. Gott KM, Potter CA, Doyle-Eisele M, Lin Y, Wilder J, Scott BR. A Comparison of Cs-137 γ Rays and 320-kV X-Rays in a Mouse Bone Marrow Transplantation Model. Dose-response : a publication of International Hormesis Society [Internet]. SAGE PublicationsSage CA: Los Angeles, CA; 2020 [cited 2022 Jan 30];18:1559325820916572. Available from: http://www.ncbi.nlm.nih.gov/pubmed/32284702

31. Epp ER, Woodard HQ, Weiss H. Energy absorption by the bone marrow of the mouse receiving whole-body irradiation with 250-Kv x-rays or cobalt-60 gamma rays. Radiation research. 1959;11:184–97.

32. Sinclair WK. The relative biological effectiveness of 22-Mevp x-rays, cobalt-60 gamma rays, and 200-Kvcp x-rays. V. Absorbed dose to the bone marrow in the rat and the mouse. Radiation research. 1962;16:369–83.

33. Plett PA, Sampson CH, Chua HL, Joshi M, Booth C, Gough A, et al. Establishing a murine model of the hematopoietic syndrome of the acute radiation syndrome. Health Physics [Internet]. 2012 [cited 2022 Jan 30];103:343–55. Available from: https://journals.lww.com/health-physics/Fulltext/2012/10000/Establishing_a_Murine_Model_of_the_Hematopoietic.3.aspx

34. Thoday JM, Vernon Hospital and M. Total X-Irradiation of Rats under Urethane Anæsthesia. Nature 1949 163:4134 [Internet]. Nature Publishing Group; 1949 [cited 2022 Feb 12];163:134–5. Available from: https://www-nature-com.elibrary.einsteinmed.edu/articles/163134a0

35. Suit HD, Sedlacek RS, Silver G, Dosoretz D. Pentobarbital Anesthesia and the Response of Tumor and Normal Tissue in the C3Hf/Sed Mouse to Radiation. Radiation Research [Internet]. Allen Press; 1985 [cited 2022 Feb 12];104:47–65. Available from: https://meridian.allenpress.com/radiation-research/article/104/1/47/37517/Pentobarbital-Anesthesia-and-the-Response-of-Tumor

36. Travis EL, Peters LJ, McNeill J, Thames HD, Karolis C. Effect of dose-rate on total body irradiation: Lethality and pathologic findings. Radiotherapy and Oncology. Elsevier; 1985;4:341–51.

37. Rotolo J, Stancevic B, Zhang J, Hua G, Fuller J, Yin X, et al. Anti-ceramide antibody prevents the radiation gastrointestinal syndrome in mice. The Journal of Clinical Investigation [Internet]. American Society for Clinical Investigation; 2012 [cited 2022 Feb 16];122:1786–90. Available from: http://www.jci.org

38. Nakamura Y, Arai F, Iwasaki H, Hosokawa K, Kobayashi I, Gomei Y, et al. Isolation and characterization of endosteal niche cell populations that regulate hematopoietic stem cells. Blood [Internet]. American Society of Hematology; 2010 [cited 2022 Feb 16];116:1422–32. Available from: https://ashpublications.org/blood/article/116/9/1422/103847/Isolation-and-characterization-of-endosteal-niche

39. Perry JM, Harandi OF, Porayette P, Hegde S, Kannan AK, Paulson RF. Maintenance of the BMP4-dependent stress erythropoiesis pathway in the murine spleen requires hedgehog signaling. Blood. Content Repository Only!; 2009;113:911–8.

40. Alamo IG, Kannan KB, Loftus TJ, Ramos H, Efron PA, Mohr AM. Severe Trauma and Chronic Stress Activates Extramedullary Erythropoiesis. The journal of trauma and acute care surgery [Internet]. NIH Public Access; 2017 [cited 2022 Jan 26];83:144. Available from: /pmc/articles/PMC5484090/

41. Rittase WB, Muir JM, Slaven JE, Bouten RM, Bylicky MA, Wilkins WL, et al. Deposition of Iron in the Bone Marrow of a Murine Model of Hematopoietic Acute Radiation Syndrome. Experimental hematology [Internet]. Exp Hematol; 2020 [cited 2022 Jan 19];84:54–66. Available from: https://pubmed.ncbi.nlm.nih.gov/32240658/

42. Chai X, Li D, Cao X, Zhang Y, Mu J, Lu W, et al. ROS-mediated iron overload injures the hematopoiesis of bone marrow by damaging hematopoietic stem/progenitor cells in mice. Scientific Reports 2015 5:1 [Internet]. Nature Publishing Group; 2015 [cited 2022 Jan 26];5:1–12. Available from: https://www.nature.com/articles/srep10181

43. Jia P, Xu YJ, Zhang ZL, Li K, Li B, Zhang W, et al. Ferric ion could facilitate osteoclast differentiation and bone resorption through the production of reactive oxygen species. Journal of Orthopaedic Research [Internet]. John Wiley & Sons, Ltd; 2012 [cited 2022 Jan 26];30:1843–52. Available from: https://onlinelibrary.wiley.com/doi/full/10.1002/jor.22133

44. Zarjou A, Jeney V, Arosio P, Poli M, Zavaczki E, Balla G, et al. Ferritin ferroxidase activity: A potent inhibitor of osteogenesis. Journal of Bone and Mineral Research [Internet]. John Wiley & Sons, Ltd; 2010 [cited 2022 Jan 26];25:164–72. Available from: https://onlinelibrary.wiley.com/doi/full/10.1359/jbmr.091002

45. Das DKR, Chakraborty A, Sinha M, Manna K, Mukherjee D, Chakraborty A, et al. Modulatory role of quercetin against gamma radiation-mediated biochemical and morphological alterations of red blood cells. http://dx.doi.org/103109/095530022013767989 [Internet]. Taylor & Francis; 2013 [cited 2022 Jan 26];89:471–81. Available from: https://www.tandfonline.com/doi/abs/10.3109/09553002.2013.767989

46. Paris F, Fuks Z, Kang A, Capodieci P, Juan G, Ehleiter D, et al. Endothelial apoptosis as the primary lesion initiating intestinal radiation damage in mice. Science [Internet]. American Association for the Advancement of Science; 2001 [cited 2022 Feb 16];293:293–7. Available from: https://www.science.org/doi/abs/10.1126/science.1060191

47. Rotolo JA, Fong CS, Bodo S, Nagesh PKB, Fuller J, Sharma T, et al. Anti-ceramide single-chain variable fragment mitigates radiation GI syndrome mortality independent of DNA repair. JCI Insight [Internet]. American Society for Clinical Investigation; 2021 [cited 2022 Feb 16];6. Available from: /pmc/articles/PMC8119204/

48. Hua G, Thin TH, Feldman R, Haimovitz-Friedman A, Clevers H, Fuks Z, et al. Crypt base columnar stem cells in small intestines of mice are radioresistant. Gastroenterology [Internet]. Gastroenterology; 2012 [cited 2022 Feb 16];143:1266–76. Available from: https://pubmed.ncbi.nlm.nih.gov/22841781/

49. Fu KK, Phillips TL, Heilbron DC, Ross G, Kane LJ. Relative Biological Effectiveness of Low- and High-LET Radiotherapy Beams for Jejunal Crypt Cell Survival at Low Doses Per Fraction1. https://doi.org/101148/1321205 [Internet]. The Radiological Society of North America ; 1979 [cited 2022 Jan 26];132:205–9. Available from: https://pubs.rsna.org/doi/abs/10.1148/132.1.205

50. Electron Spectra and the RBE of X Rays on JSTOR [Internet]. [cited 2022 Feb 16]. Available from: https://www.jstor.org/stable/3580692?seq=1#metadata_info_tab_contents

51. Saha S, Bhanja P, Kabarriti R, Liu L, Alfieri AA, Guha C. Bone Marrow Stromal Cell Transplantation Mitigates Radiation-Induced Gastrointestinal Syndrome in Mice. PLOS ONE [Internet]. Public Library of Science; 2011 [cited 2022 Feb 3];6:e24072. Available from: https://journals.plos.org/plosone/article?id=10.1371/journal.pone.0024072

52. Storer JB, Harris PS, Furchner JE, Langham WH. The Relative Biological Effectiveness of Various Ionizing Radiations in Mammalian Systems. Radiation Research. JSTOR; 1957;6:188.

