## Supplementary Methods for "Increased Relative Biological Effectiveness of Orthovoltage X-rays Compared to γ-rays in Preclinical Irradiation"

### **<u>Supplemental Methods:</u>**

#### Spleen Flow Cytometry Panel

| Marker | Conjugate | Dilution | Company | Catalog | Clone |
|---|---|---|---|---|---|
| CD11b | BUV395 | 1:1000 | BD | 563553 | M1/70 |
| CD19 | BUV661 | 1:1000 | BD | 612971 | 1D3 |
| NKp46 | BUV737 | 1:50 | BD | 612805 | 29A1.4 |
| CD8 | BUV805 | 1:500 | BD | 612898 | 53-6.7 |
| CD4 | BV510 | 1:200 | BD | 563106 | RM4.5 |
| F4/80 | BV650 | 1:300 | Biolegend | 123149 | BM8 |
| CD11c | BV750 | 1:400 | Biolegend | 117357 | N418 |
| B220 | FITC | 1:50 | eBioscience | 11-0452-82 | RA3-6B2 |
| CD45 | AF532 | 1:100 | eBioscience | 58-0451-82 | 30-F11 |
| CD3ε | PerCP | 1:100 | Biolegend | 100326 | 145-2C11 |
| CD49b | PE | 1:100 | Biolegend | 108907 | DX5 |
| Ly-6G | APC | 1:400 | Biolegend | 127614 | 1A8 |
| MHC II | AF700 | 1:200 | Biolegend | 107622 | M5/114.15.2 |
| LD | Zombie NIR | 1:1500 | | | |

#### HSPC Flow Cytometry Panel

| Marker | Conjugate | Dilution | Company | Catalog | Clone |
|---|---|---|---|---|---|
| Anti-Mouse Lineage Cocktail | Biotin | 1:200 ea | Biolegend | 79752 | |
| Sca-1 | AF700 | 1:100 | Biolegend | 108142 | D7 |
| c-Kit | APC | 1:100 | Biolegend | 105811 | 2B8 |
| CD135 | PE | 1:100 | Biolegend | 135305 | A2F10 |
| CD34 | BV421 | 1:50 | Biolegend | 152208 | SA376A4 |
| CD150 | BV711 | 1:100 | Biolegend | 115941 | TC15-12F12.2 |
| CD48 | BUV395 | 1:200 | BD | 740236 | HM48-1 |
| CD16/32 | BV480 | 1:200 | BD | 746324 | 2.4G2 |
| CD127 | BV650 | 1:100 | Biolegend | 135043 | A7R34 |
| Streptavidin | BV570 | 1:100 | Biolegend | 405227 | |
| LD | Zombie NIR | 1:1500 | | | |
