## Supplementary Figures for "Increased Relative Biological Effectiveness of Orthovoltage X-rays Compared to γ-rays in Preclinical Irradiation"

#### Supplemental Figure 1

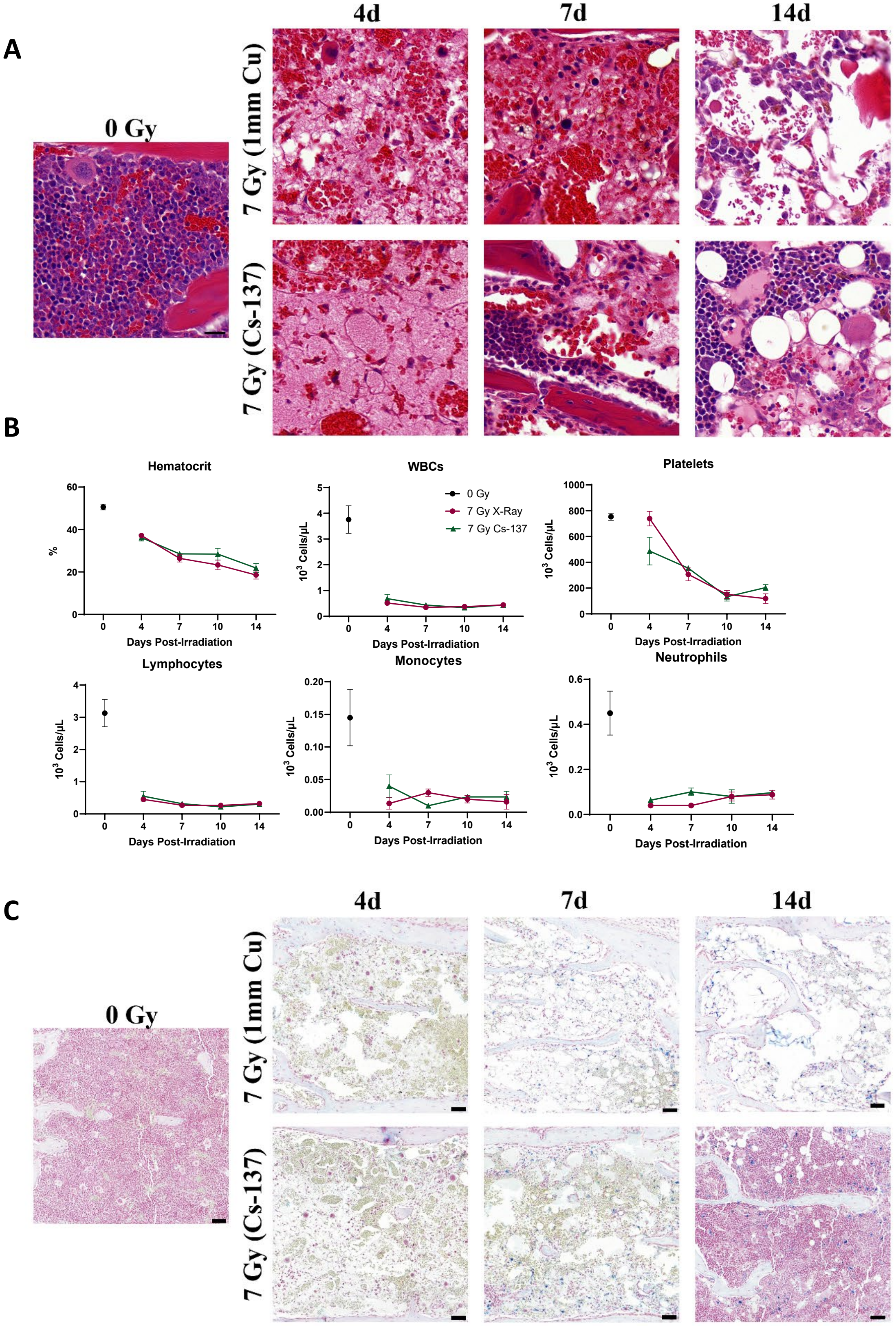


**Supplemental Figure 1:** (A) Representative H&E-stained sternal bone marrow from 4-14 days post-irradiation. (B) Complete blood count parameters from 4-14 days post-irradiation. (C) Prussian Blue-stained sternal bone marrow from 4-14 days post-irradiation.

#### Supplemental Figure 2

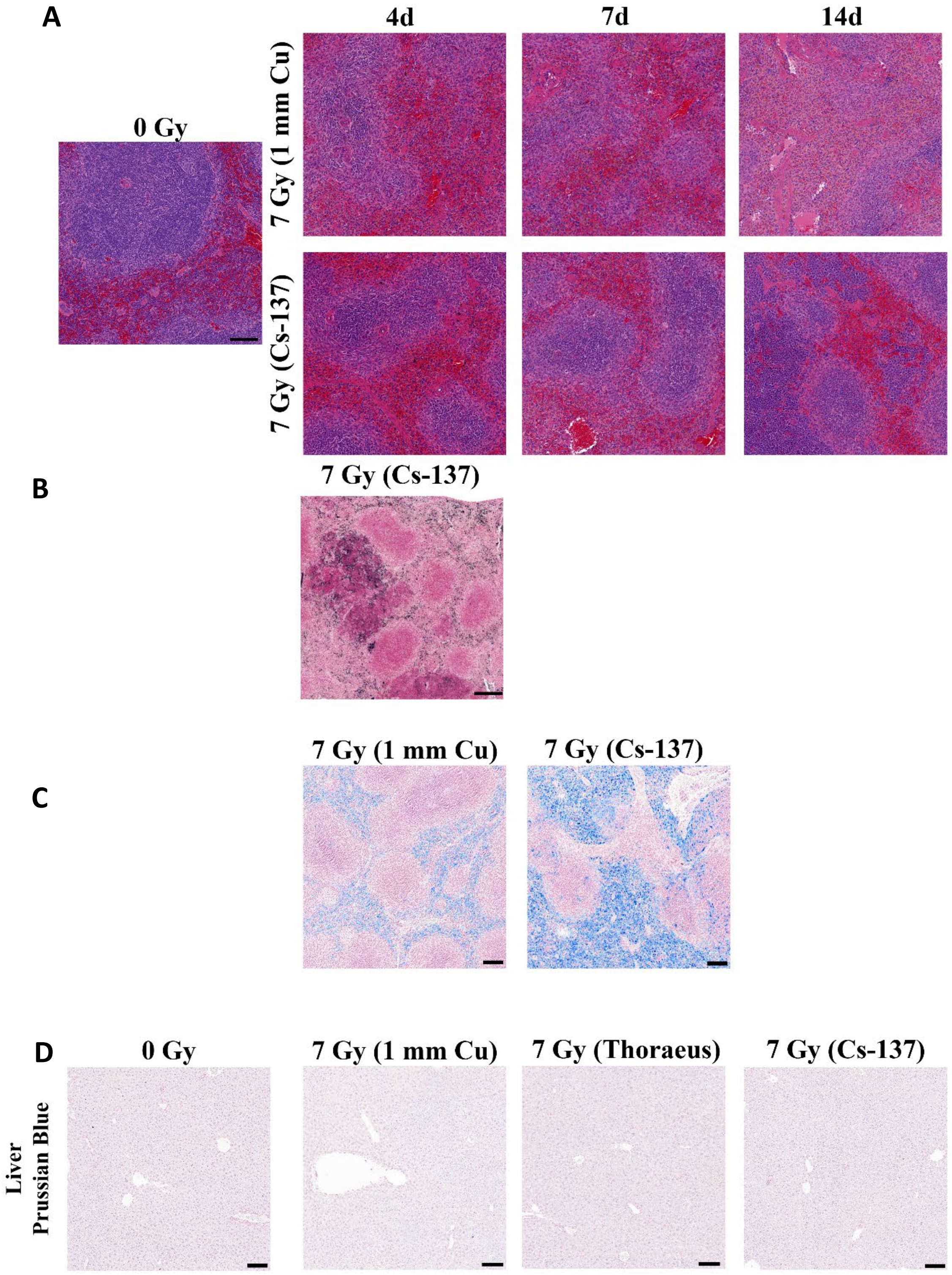


**Supplemental Figure 2:** (A) Representative H&E-stained spleens from 4-14 days post-irradiation. (B) Representative myeloperoxidase immunohistochemistry in splenic nodules 10 days post-irradiation with $^{137}$Cs. (C) Representative Prussian Blue-stained spleens 14 days post-irradiation. (D) Prussian Blue-stained livers 10 days post-irradiation.

### Supplemental Figure 3

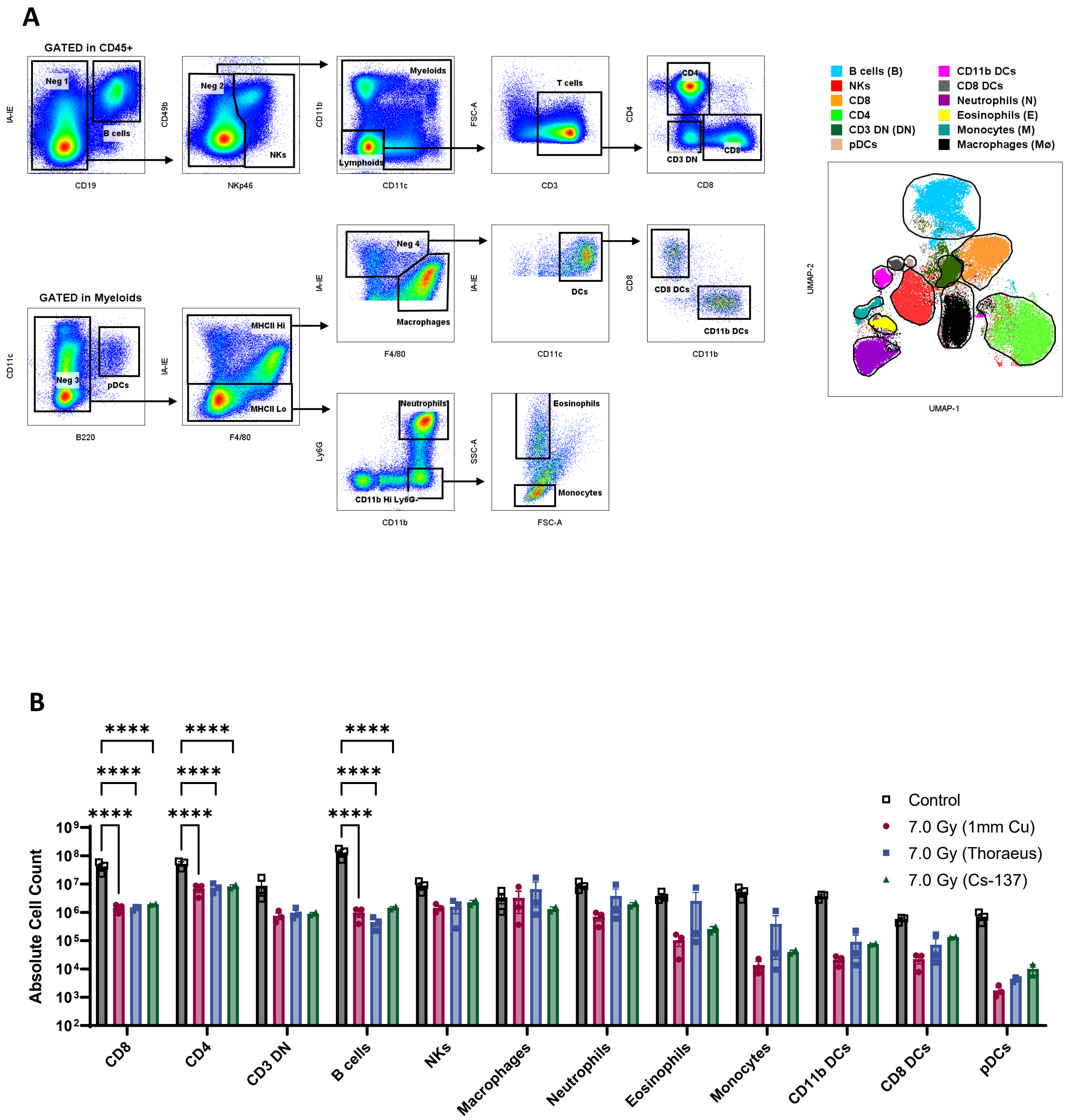


**Supplemental Figure 3:** (A) Manual Gating strategy used to validate UMAP clusters. Manual gating was overlaid over the UMAP clustering at right. (B) Absolute counts of splenic immune populations 10-days post-irradiation.

#### Supplemental Figure 4

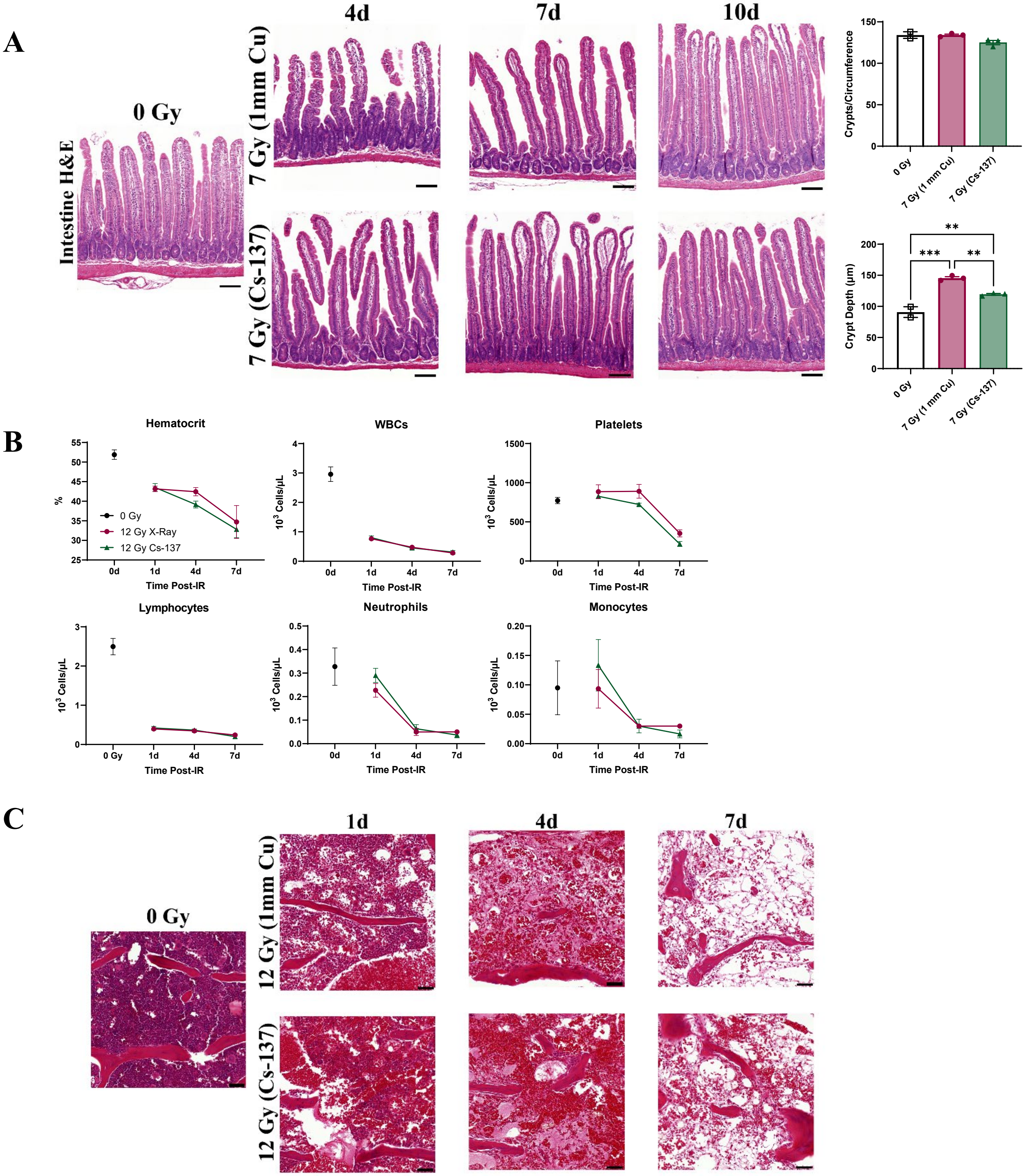


**Supplemental Figure 4:** (A) Representative H&E-stained intestines from 4-10 days post-irradiation with associated measurements of crypt depth and crypt loss. (B) Complete blood count parameters from 1-7 days post-irradiation. (C) Representative H&E-stained sternal bone marrow from 1-7 days post-irradiation

#### Supplemental Figure 5

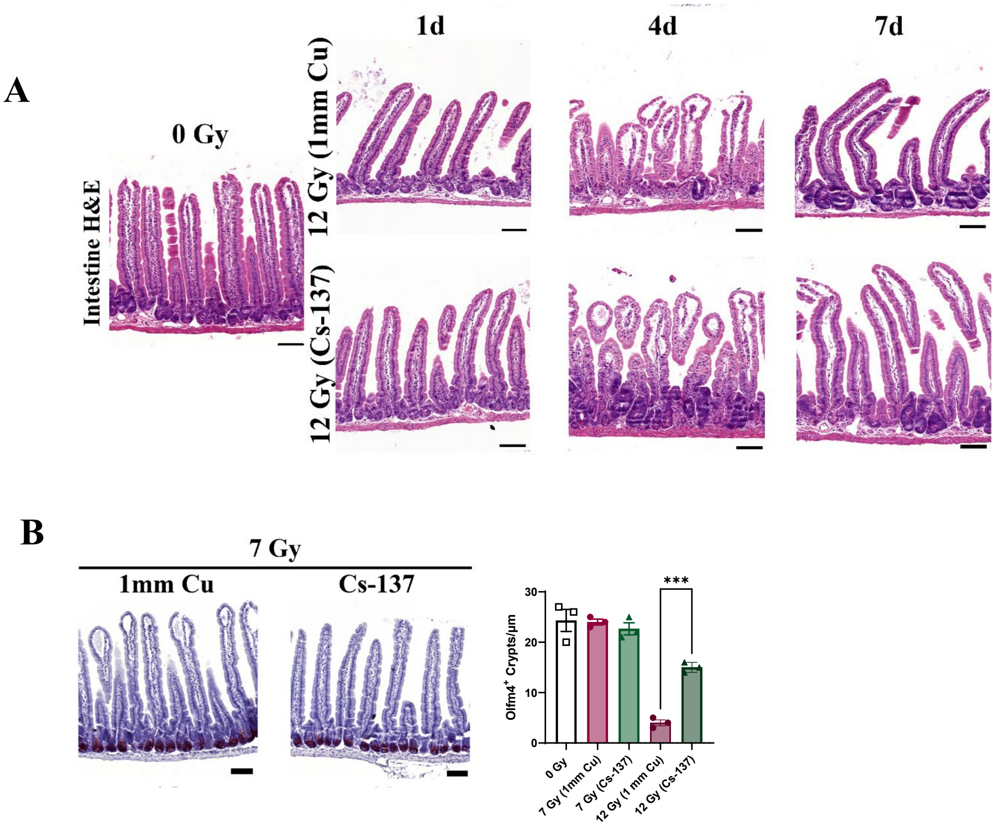


**<u>Supplemental Figure 5:</u>** (A) Representative H&E-stained intestines from 1-7 days post-irradiation. (B) Olfm4 IHC 4 days post-7 Gy WBI with associated counts per micron.
